# NaDyNet: A Toolbox for Dynamic Network Analysis of Naturalistic Stimuli

**DOI:** 10.1101/2024.11.29.626037

**Authors:** Junjie Yang, Zhe Hu, Junjing Li, Xiaolin Guo, Xiaowei Gao, Jiaxuan Liu, Yaling Wang, Zhiheng Qu, Wanchun Li, Zhongqi Li, Wanjing Li, Yien Huang, Jiali Chen, Hao Wen, Binke Yuan

**Author notes:** Correspondence: Binke Yuan, PhD., Room 206, Institute for Brain Research and Rehabilitation, South China Normal University, Guangzhou, China Postcode: 510631.

## Abstract

Experiments with naturalistic stimuli (e.g., listening to stories or watching movies) are emerging paradigms in brain function research. The content of naturalistic stimuli is rich and continuous. The fMRI signals of naturalistic stimuli are complex and include different components. A major challenge is isolate the stimuli-induced signals while simultaneously tracking the brain’s responses to these stimuli in real-time. To this end, we have developed a user-friendly graphical interface toolbox called NaDyNet (Naturalistic Dynamic Network Toolbox), which integrates existing dynamic brain network analysis methods and their enhanced versions. The main features of NaDyNet are: 1) extracting signals of interest from naturalistic fMRI signals; 2) incorporating six commonly used dynamic analysis methods and three static analysis methods; 3) enhanced versions of these dynamic methods by adopting inter-subject analysis to eliminate the effects of non-interest signals; 4) performing K-means clustering analysis to identify temporally reoccurring states along with their temporal and spatial attributes. We then introduced the rationale for incorporating inter-subject analysis to improve existing dynamic brain network analysis methods, and presented numerous examples. We also summarized research progress in comparing methodological efficacy, offered our recommendations for method selection in dynamic network analysis, and discussed the limitations of current approaches and directions for future research. We hope that this open source toolbox will promote the development of naturalistic neuroscience. The toolbox is available at https://github.com/yuanbinke/Naturalistic-Dynamic-Network-Toolbox.

## 1 Introduction

A fundamental goal of cognitive neuroscience is to elucidate the mechanisms through which the brain interacts with the external environment. To achieve this, scientists have developed numerous paradigms over the past decades, including block, event-related, and resting-state paradigms (Finn, 2021). One of the main limitations of these paradigms is their low ecological validity (Cole, Smith, & Beckmann, 2010; Lurie et al., 2020). Naturalistic paradigms, such as listening to stories or watching movies, have recently received increasing attention from researchers (Jaaskelainena, Sams, Glerean, & Ahveninen, 2021; Saarimaki, 2021; Sonkusare, Breakspear, & Guo, 2019; Willems, Nastase, & Milivojevic, 2020). Naturalistic stimuli approximate real-life experiences, are more engaging, and are relatively easy to implement. The features of naturalistic stimuli are rich, including words, sentences, narratives, music, social interactions, actions, etc. In addition, these features have their own dimensions (e.g., semantic dimensions for words) (Binder et al., 2016; S. Wang et al., 2023) and attributes (e.g., positive, negative, and neutral) (Kousta, Vinson, & Vigliocco, 2009; Vine, Boyd, & Pennebaker, 2020). Therefore, naturalistic stimuli has high ecological validity (Finn, Glerean, Hasson, & Vanderwal, 2022), test-retest reliability (Gao et al., 2020; J. Wang et al., 2017), and can be used to study different aspects (Jaaskelainena et al., 2021), such as language processing (Hamilton & Huth, 2020; Huth, de Heer, Griffiths, Theunissen, & Gallant, 2016; Zada et al., 2024), memory construction (Chien & Honey, 2020), attention engagement (Song, Finn, & Rosenberg, 2021), social communication (Yeshurun, Nguyen, & Hasson, 2021), and emotion recognition (Lin, Zhang, Wang, & Wang, 2024; Saarimaki, 2021). Various studies also adopted naturalistic stimuli for clinical populations (Eickhoff, Milham, & Vanderwal, 2020; Zuo et al., 2020), such as ADHD (Salmi et al., 2020), schizophrenia (Yang et al., 2020), autism spectrum disorder (Bolton, Freitas, Jochaut, Giraud, & Van De Ville, 2020), depression (Gruskin, Rosenberg, & Holmes, 2020), and glioma for presurgical mapping (Yao et al., 2022).

The richness of naturalistic stimuli also poses challenges for analytical methods. Existing analytical methods, such as inter-subject correlation (Nastase, Gazzola, Hasson, & Keysers, 2019) and inter-subject functional connectivity (Simony et al., 2016), still focus on "static" activation and functional connectivity (SFC). However, the nature of naturalistic stimuli is dynamic and real-time. Therefore, a major challenge in adopting naturalistic stimuli is how to isolate stimulus-induced signals and track the brain’s corresponding responses in real time. In the field of resting-state fMRI, numerous dynamic brain network analysis methods have been developed (Torabi, Mitsis, & Poline, 2024; Xie et al., 2019; Binke Yuan et al., 2024), such as flexible least squares (FLS) (Kalaba & Tesfatsion, 1989; Liao et al., 2014), dynamic conditional correlation (DCC) (Lindquist, Xu, Nebel, & Caffo, 2014; B. Yuan, Xie, Gong, et al., 2023; B. Yuan, Xie, Wang, et al., 2023), general linear Kalman filter (GLKF) (Pascucci, Rubega, & Plomp, 2020), multiplication of temporal derivatives (MTD) (Shine et al., 2015), sliding-window functional connectivity with L1-regularization (SWFC) (Allen et al., 2014), hidden Markov models (HMM) (K. Chen et al., 2022), hidden semi-Markov models (HSMM) (Shappell, Caffo, Pekar, & Lindquist, 2019), Co-activation pattern (Liu & Duyn, 2013; Liu, Zhang, Chang, & Duyn, 2018), and Mapper (Hasegan, Geniesse, Chowdhury, & Saggar, 2023; Saggar, Shine, Liegeois, Dosenbach, & Fair, 2022; Zhang, Chowdhury, & Saggar, 2023), etc. These methods have demonstrated their accuracy and reliability for resting-state fMRI signals (Choe et al., 2017; Lindquist et al., 2014; Shappell et al., 2019; Torabi et al., 2024; Xie et al., 2019), but they cannot be directly applied to naturalistic stimuli.

For naturalistic stimuli, the components in the measured BOLD (blood oxygen level-dependent) signals are complex. In simple terms, naturalistic signals can be seen as a combination of three components: 1) stimulus-induced signals; 2) intrinsic neural signals; and 3) non-neural signals (such as head movement and respiration) (Nastase et al., 2019; Simony et al., 2016). The first component is shared across subjects, while the latter two are subject-specific and uncorrelated. A straightforward approach to component separation is the introduction of ISC analysis (Hasson, Nir, Levy, Fuhrmann, & Malach, 2004). In our recent study, we demonstrated that combining above methods with ISC (i.e., ISFLS, ISDCC, ISGLKF, ISMTD, ISSWFC, ISHMM, and ISHSMM) can effectively isolate stimulus-induced components, resulting in robust, consistent network patterns across subjects (L. Chen et al., 2024; Binke Yuan et al., 2024). We also systematically compared the efficacy of these methods using simulated data. We found that all the enhanced methods outperformed their original versions in tracking transient network reorganization (Binke Yuan et al., 2024). However, most of these algorithms require code debugging for implementation and lack user-friendly graphical interfaces.

To make these methods more accessible and to advance the field of naturalistic neuroscience, we have developed a user-friendly toolbox called NaDyNet (Naturalistic Dynamic Network Toolbox). This toolbox supports the extraction of signals of interest and the construction of dynamic networks. It provides six dynamic network analysis methods (FLS, DCC, GLKF, MTD, SWFC, CAP), their enhanced versions (ISFLS, ISDCC, ISGLKF, ISMTD, ISSWFC, ISCAP), and three static network analysis methods (SFC, ISC, ISFC). It also includes a subsequent K-means clustering analysis to determine temporally reoccurring states and their state transition properties, with a visualization interface. To our knowledge, this is the most comprehensive dynamic brain network analysis toolbox available to date.

We would like to begin by expressing our deep respect for the researchers who developed these methods. We are also sincerely grateful for their decision to make these methods open source, allowing us to create a graphical interface for them. In the following sections, we first introduced the modules in NaDyNet and demonstrated how to use our toolbox effectively. Next, we gave a brief overview of the mathematical principles underlying these methods, along with a discussion of the free parameters and inherent limitations associated with each metric. We then explained the logic of the ISC and provided examples that highlight the advantages of the enhanced versions using the ISC. We also summarized recent research progress on the methodological efficacy and reliability of these metrics. Finally, in the discussion section, we offered our recommendations on how to select the most appropriate metrics for your study, as well as insights into the future of dynamic network analysis.

## 2 Functionality of NaDyNet

NaDyNet is an open source and free toolkit developed in MATLAB 2018a and licensed under the General Public License (GPL). NaDyNet is compatible with several versions of MATLAB, including 2018b, 2019a, 2019b, 2020a, 2020b, 2021a, 2021b, 2022a, and 2022b. Depending on whether the analysis methods are dynamic in nature, the software classifies them into two primary categories: dynamic brain network analysis methods and static brain network analysis methods. Dynamic brain network analysis methods include FLS, SWFC, DCC, MTD, GLKF, CAP, and their enhanced versions: ISFLS, ISSWFC, ISDCC, ISMTD, ISGLKF, and ISCAP. Static brain network analysis methods include SFC, ISC, and ISFC. In addition, these methods can be further divided into two categories depending on whether the analysis approach is ROI-wise or voxel-wise: ISC, CAP, and ISCAP are voxel-wise analysis methods, while the remaining metrics are ROI-wise.

The main interface of this MATLAB toolbox consists of three modules: the ROI Time Course Extraction Module, the Method Selection Module, and the Clustering and Plotting Module, as shown in **Figure 1**. (i) The ROI Time Course Extraction Module primarily extracts the time series for each ROI to facilitate ROI-wise analysis. (ii) Within the Method Selection Module, users can select from 11 ROI-wise and 3 voxel-wise methods according to their specific needs. Once a method is selected, the corresponding graphical user interface (GUI) is launched. Users must then specify path to the data required for the brain network analysis, configure the method’s free parameters (as outlined in Table 1), and proceed to perform the analysis. (iii) The Clustering and Plotting Module allows users to perform K-means clustering on the dynamic ROI-wise brain network analysis outcomes. This involves estimating the optimal number of clusters (K) and generating both the connectivity matrices and state transition matrices for each state associated with the optimal K. We then provide a concise description of the steps users need to take, along with the analytical options available at each state of the process.

**Figure 1.**
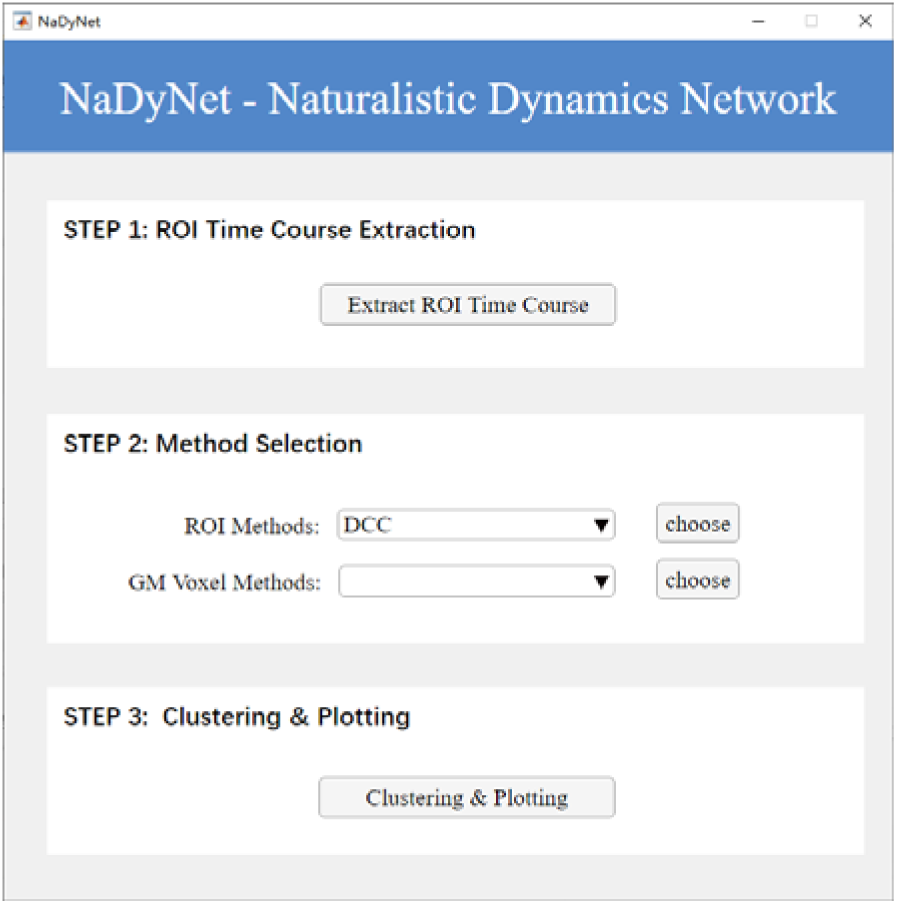
Graphical user interface of NaDyNet. Three modules (steps) present in the main interface.

**Table 1.**
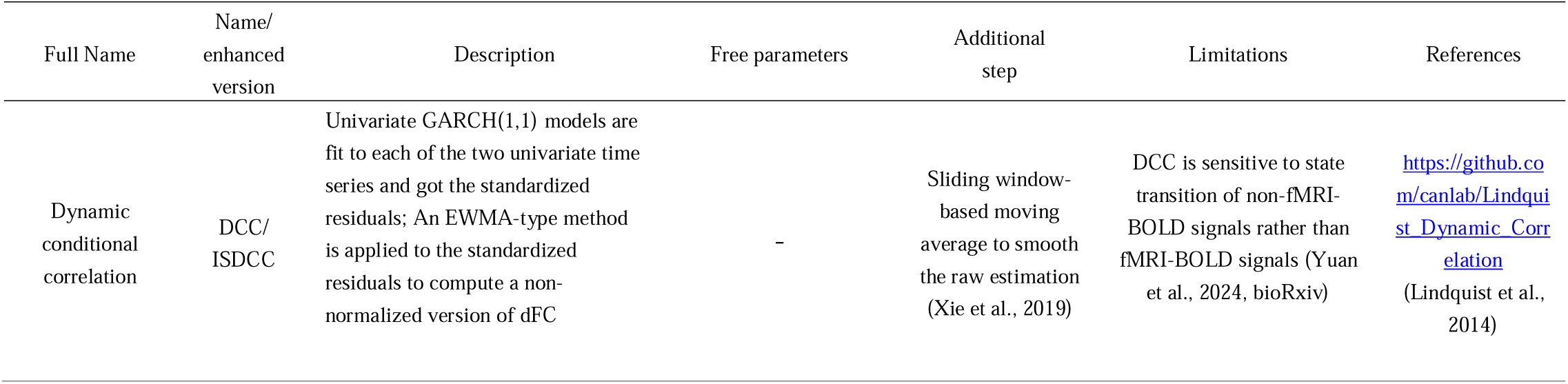
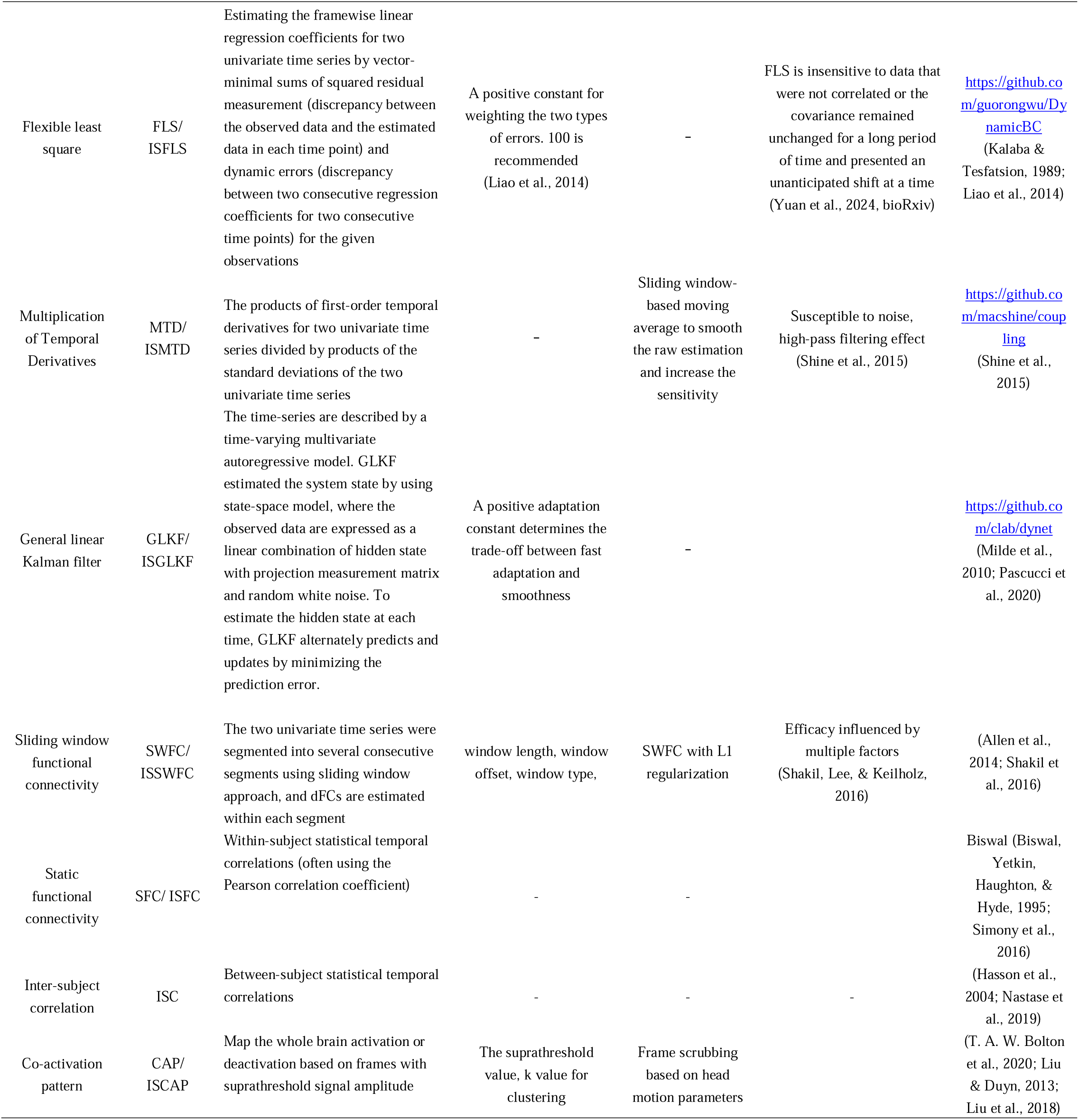
Summary of methods available in NaDyNet

### 2.1 ROI Time Course Extraction

The interface of the ROI Time Course Extraction Module is shown in **Figure 2**. This module facilitates the extraction of ROI time courses from fMRI data for a group of subjects. If voxel-wise methods are used, this step is not necessary. Before performing this step, it is important to ensure that each subject’s fMRI data have been pre- processed (Chao-Gan & Yu-Feng, 2010; Jia et al., 2019; J. Wang et al., 2015; Yan, Wang, Zuo, & Zang, 2016) and been registered in MNI space, aligned with the ROI mask. Similar to other types of fMRI data, there is no standardized preprocessing pipeline for fMRI data with naturalistic stimuli. Based on a review of several high-impact studies (Chien & Honey, 2020; Song, Finn, et al., 2021; Song, Park, Park, & Shim, 2021), we recommend at least the following preprocessing steps: (1) motion correction; (2) registration to standard space; (3) detrending; and (4) Gaussian smoothing. Since dynamic brain network analysis methods are particularly sensitive to noise and motion artifacts, we also recommend the following additional steps: (1) regression of motion, white matter, and CSF signals; and (2) low-pass filtering between 0.01-0.1 Hz. Given the ongoing debate around global signal regression (Murphy & Fox, 2017), we do not make a formal recommendation for its use in this context.

**Figure 2.**
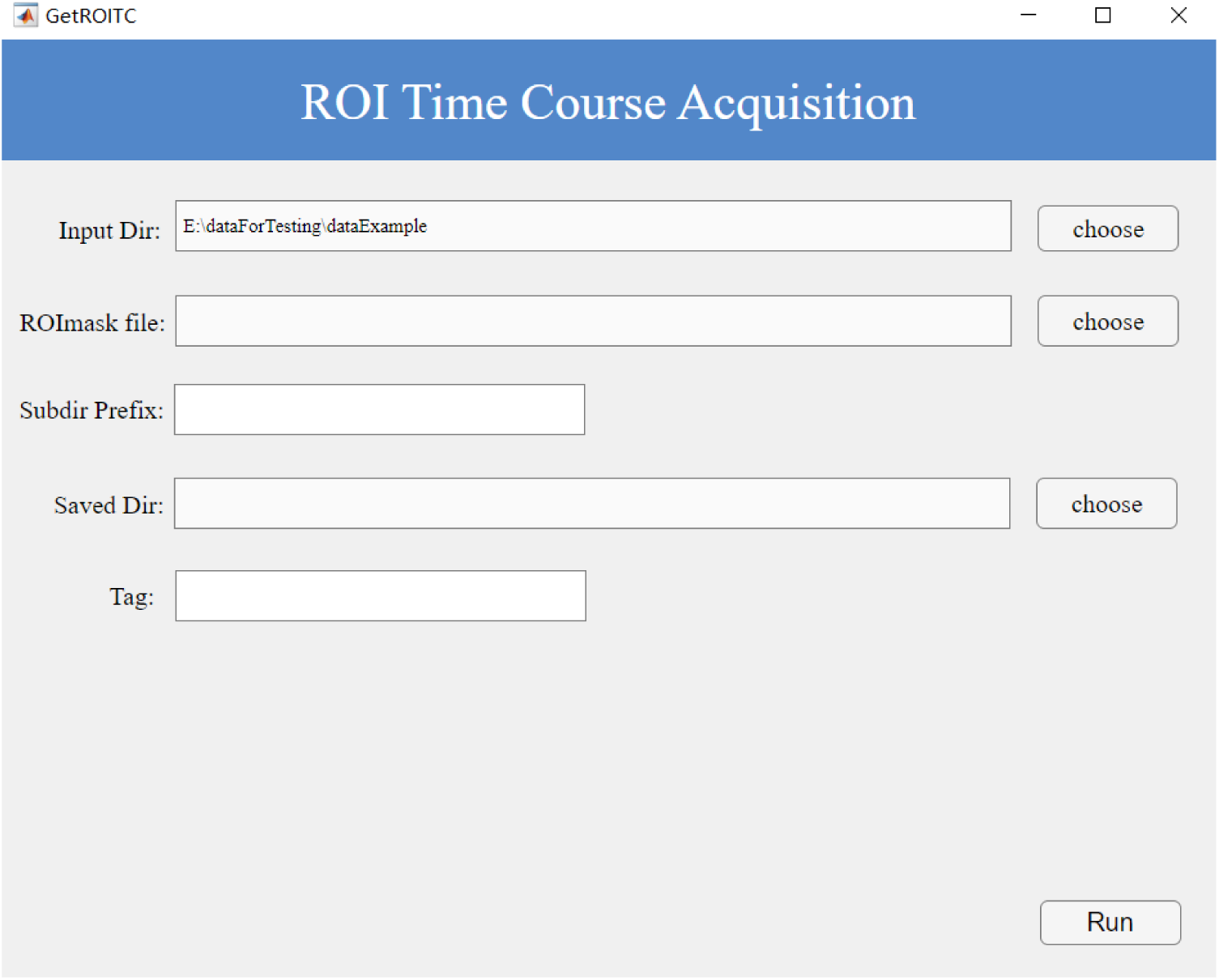
The ROI time course extraction interface

Prior to data extraction, each participant’s dataset must be organized according to certain conventions: (i) the 4D fMRI file for each participant should be stored in an individual subfolder; (ii) each subfolder should follow a systematic naming convention, including a consistent prefix. As shown in **Figure 3**, in the "dataExample2" directory, the folders for three participants could be named as sub- UTS01, sub-UTS02, and sub-UTS03, all prefixed with ’sub’ to facilitate selection within the toolbox. Having organized the data for each subject according to this structured framework, the user can proceed to operate the module by following the steps below.

**Figure 3.**
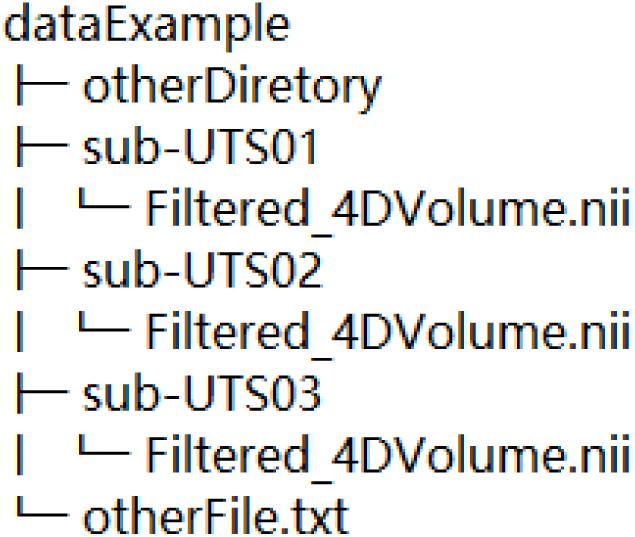
Organizational structure of NifTi files for ROI-wise methods.

#### Operational steps

1. Select the pre-processed fMRI data folder that meets the above criteria as the input path.
2. Select an ROI mask file. By default, each ROI within this mask file is assigned a unique identifier, and the number of identifiers must be equal to the number of ROIs. Non-sequential numbering may result in the presence of null data. It is essential that the voxel dimensions and shape of the ROI mask file match those of the participant’s fMRI data; otherwise, errors will occur.
3. As the folders for our participants are systematically named with a common prefix, ’sub’, simply entering the correct prefix will prompt the interface to display the number of identified participants.
4. Specify an output path to store the corresponding results.
5. Given the variety of ROI mask types, please enter a distinguishable label tag to avoid unnecessary confusion in subsequent analyses.

The toolbox processes each participant sequentially, reading their 4D NifTi files and extracting the time series signals of all voxels for each region based on the coordinates specified in the ROI mask. It then calculates the average signal value for each region. Finally, a corresponding MAT file is generated for each participant in the specified output path, with the data structure of each MAT file formatted as nT×nROI. (Ultimately, two corresponding MAT files will be generated for each subject in the output path—one stored in the ’raw’ folder and the other in the ’zscore’ folder, with the latter containing data that has undergone z-score normalization. The data structure within each MAT file is formatted as *nT*×*nROI*, where *nT* denotes the number of time points, and *nROI* represents the number of regions of interest, and the same applies throughout). Users are free to choose any format of the data based on their needs for subsequent analyses.

### 2.2 Method Selection

In brain network analysis, the selection of the appropriate method is crucial, as different approaches can reveal either dynamic or static features of brain networks, thereby influencing the interpretation of the results. This module, the Method Selection Module, is designed to provide the user with a range of methods for brain network analysis, covering both ROI-wise and voxel-wise techniques (**Figure 4**). Users have the flexibility to choose the most appropriate method based on specific research questions and the characteristics of their data. By making an informed choice of method, users can effectively analyze complex patterns of brain activity, leading to more reliable results in subsequent analyses.

**Figure 4.**
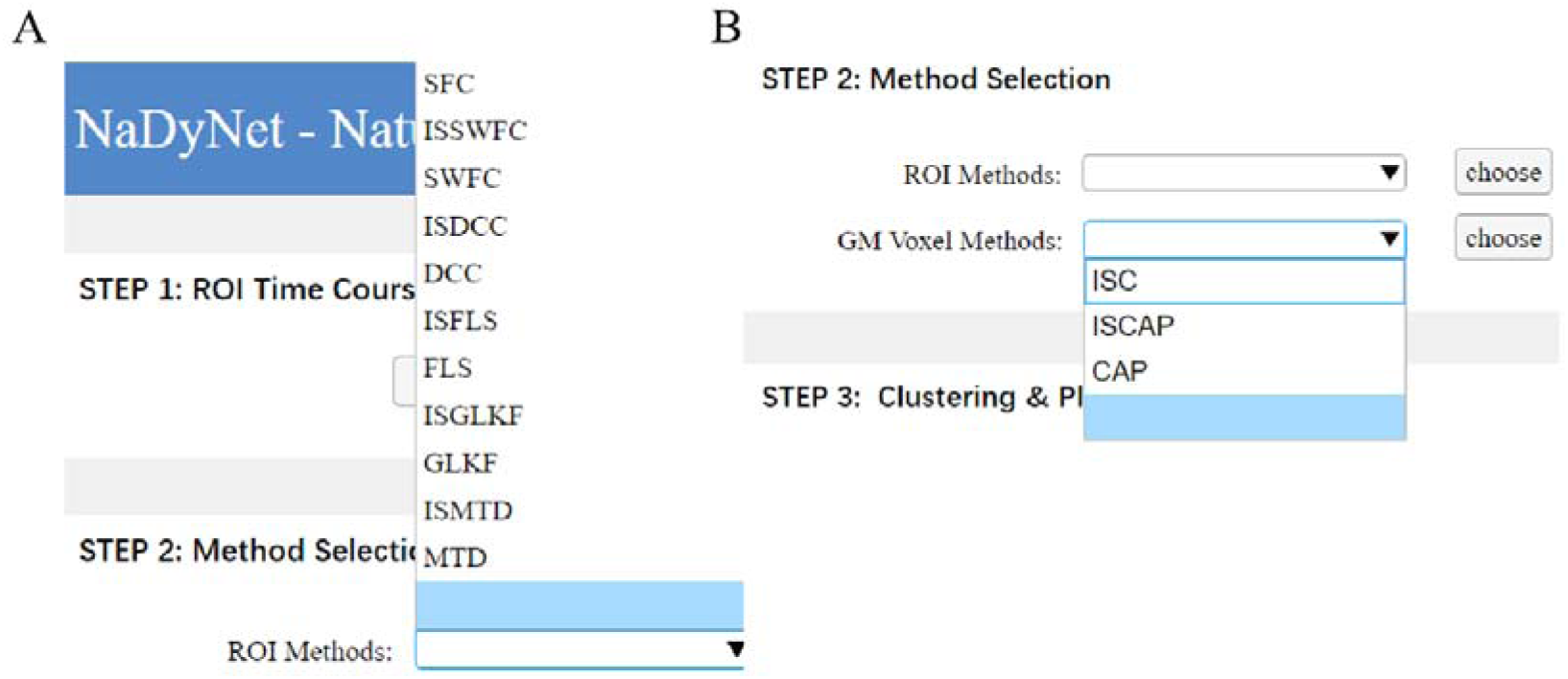
The method selection interface. Eleven ROI methods and three voxel-wise methods appear in the drop-down lists.

#### 2.2.1 ROI-wise Method

The ROI-wise approach is a technique that focuses on pre-defined brain regions, known as regions of interest (ROIs), for analysis. This method divides the brain into multiple regions that are either functionally or anatomically significant (Fan et al., 2016; Glasser et al., 2016; Yeo et al., 2011), allowing researchers to study the functional connectivity or activity patterns between these areas. Each ROI represents a region of interest, typically defined by standardized brain templates (e.g., AAL, Brainnetome) or generated according to specific experimental needs. The core principle of ROI-wise analysis is to aggregate the time signals of all voxels within each ROI to calculate the average signal for that region. This approach reduces the dimensionality of the data and increases the signal-to-noise ratio (SNR).

A total of 11 ROI-wise methods are presented in this section, with a brief description of the principles underlying each method given in Table 1. Due to space limitations, the mathematical formulae for each method can be found in the corresponding references. In Yuan et al. (2024, bioRxiv), we summarized the mathematical formulations for most of these methods. It is noteworthy that the calculation of SFC is based on the Pearson correlation coefficient, which is widely recognized as one of the most commonly used calculation methods. Apart from SFC, the remaining ten ROI analysis methods are dynamic analysis techniques, namely FLS, SWFC, DCC, MTD, GLKF, and CAP and their enhanced versions: ISFLS, ISSWFC, ISDCC, ISMTD, and ISGLKF. We offer two improved enhancement methods; both methods are applicable to ROI-wise analyses, while the second method is also applicable to voxel-wise analyses.

##### (1) Leave-One-Out Average Approach (LOO)

For each subject, the original data (dimensions: [*nT, nROI*], where *nT* is the number of time points and *nROI* is the number of regions of interest) is left out. The average group data, LOO-mean (dimensions: [*nT, nROI*]), is then calculated using the remaining subjects. The subject’s time course (TC) data and the inter-subject consistent data, LOO-Average, are merged into a new matrix (dimensions: [*nT, 2nROI*]), based on which a dynamic brain network matrix is computed, with dimensions [*2nROI*, *2nROI*, (*nT - wsize*)], where *wsize* is the window size. For framewise methods, *wsize* is set to 0. In this dynamic brain network matrix, the top-left and bottom-right quadrants represent the dynamic connectivity between the original data, while the top-right and bottom-left quadrants capture the dynamic connectivity between subjects, i.e., the enhanced version of the result. In our recent work, we used this approach to compare the performance of DCC and ISDCC and demonstrated its effectiveness (L. Chen et al., 2024).

##### (2) Leave-One-Out Average Regress Approach (LOO + Regress)

For a given subject, the LOO-Average data are first used to estimate the task-evoked mean (dimensions: [*nT, nROI*]). A linear regression is then performed, with the estimated task-evoked mean as the independent variable and the subject’s original signal as the dependent variable. This gives the regression coefficient (b) and the residual (*r*). Subtracting the residual (*r*) from the original signal effectively removes task-unrelated signals, leaving behind only the task-evoked signal.

The user can select any method, after which a new interface will appear allowing the user to select the folder containing the ROI time course (TC) results generated in Step 1 as the input for the ROI-wise method. It is important to note that this folder should only contain MAT files related to the ROI TC; unrelated MAT files should not be present in this location. After specifying the output folder and configuring the free parameters (as detailed in Table 1), users can simply click the button to automatically start the analysis. The 11 ROI-wise methods include 10 dynamic analysis techniques. In addition to producing individual dynamic brain network analysis results for each participant, these methods also produce a consolidated output that includes the above analysis results as well as relevant information required for visualization in the clustering and plotting module. The filename of this output ends with "_all.mat" and serves as input to the module.

#### 2.2.2 Voxel-wise methods

Voxel-wise analysis is characterized by its focus on individual signal variations at the level of each voxel (the smallest distinguishable unit) in fMRI images. This approach provides greater spatial resolution than ROI-wise methods. Voxel-wise analysis preserves the unique activity patterns of each voxel, rather than averaging signals within predefined brain regions, providing a more detailed and localized view of brain dynamics. This is particularly useful for studies that aim to capture subtle activity patterns, small neural responses or localized functional connectivity that may be lost when signals are aggregated across larger brain areas. This toolbox provides implementations of voxel-wise methods, including ISC, CAP, and an enhanced version known as ISCAP, which is applied with the Leave-One-Out Average Regress Approach.

For CAP and ISCAP analyses, we recommend the data organization structure shown in **Figure 5**. In the "dataExample2" directory, each participant’s 4D fMRI file and corresponding motion file should be stored in a separate folder. The motion file should have a .txt extension. In addition, folder names should follow a consistent naming convention, such as starting with the prefix ’sub’. This approach ensures efficient selection of the required data while excluding irrelevant files and folders. As CAP and ISCAP are frame-based analysis metrics, the motion files are used to exclude time points with excessive head movement. A threshold for excessive motion must therefore be defined. Frame-wise displacement is calculated based on the motion file, with 0.5 being a commonly used threshold. If no motion file is provided, the software defaults to a motion value of zero, meaning that the calculation of CAP and ISCAP takes into account data from all time points. It is important to note that the duration of the motion file must match that of the fMRI file, otherwise an error will occur. In addition to these metrics, several other parameters need to be configured, as detailed in Table 1.

**Figure 5.**
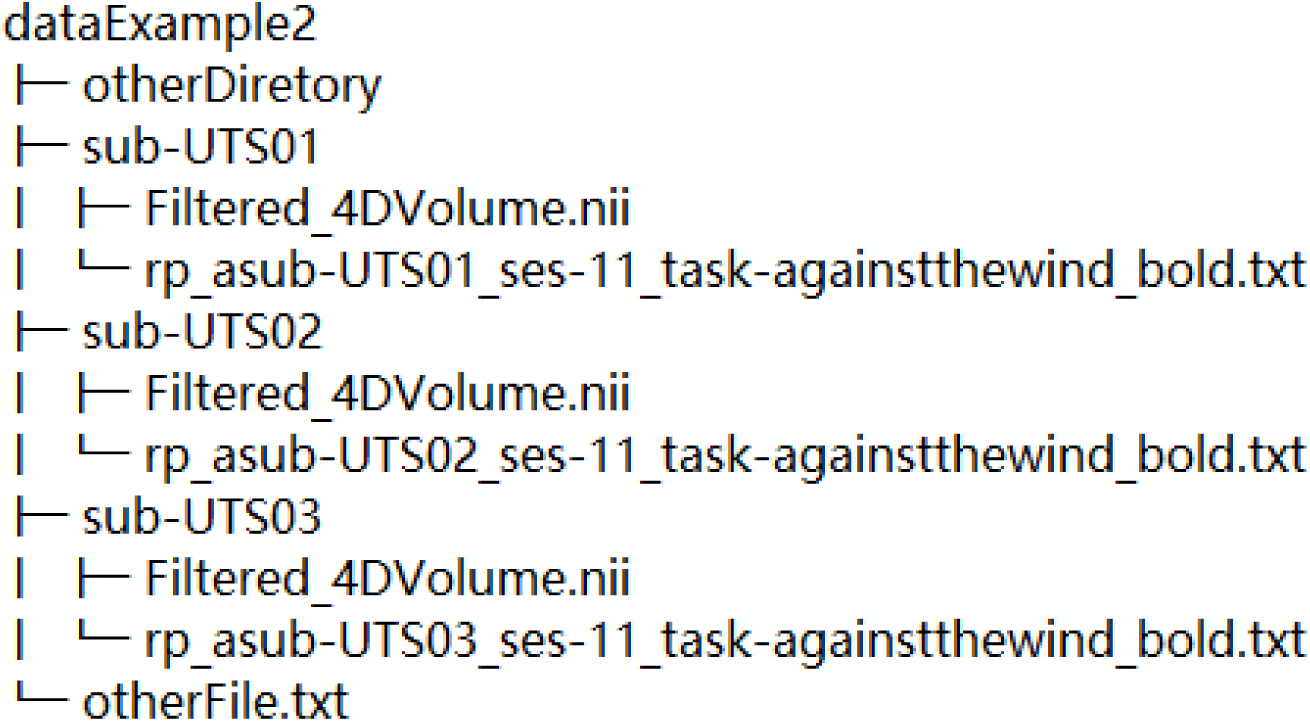
Organizational structure of files for voxel-wise methods

For the ISC method, there is no need to use head motion files, and the file organization should be structured as shown in **Figure 5**. Only the path of the pre- processed fMRI data needs to be provided. The ISC results include not only the original Pearson correlation coefficients, but also the Fisher Z-transformed results. Based on individual ISC results, mean ISC values across subjects can be calculated to determine the magnitude and intensity of task-induced activation, or they can be subjected to statistical tests such as one-sample t-tests.

#### 2.2.3 Clustering & Plotting

In brain network analysis, the Clustering and Plotting module is a critical component designed to help researchers effectively interpret and visualise complex patterns of brain activity. The primary function of clustering analysis is to group brain network data across multiple time points or samples to identify similar patterns of activity and potential functional connections. This approach not only reveals interactions between different brain regions, but also helps to understand dynamic changes in brain networks across different states. Through clustering analysis, researchers can extract meaningful insights from large-scale neuroimaging data, identify specific functional brain regions and explore their role in cognitive tasks or emotional responses. In addition, this module supports visualisation of results, allowing researchers to intuitively understand clustering results and their relationship to brain network structures. By providing high-quality graphical representations, researchers can better interpret their data and effectively communicate their findings in academic settings.

The interface of the Clustering and Plotting module is divided into three sections: determining the optimal K-value, performing the K-means clustering analysis and visualizing the results, as shown in **Figure 6**.

**Figure 6.**
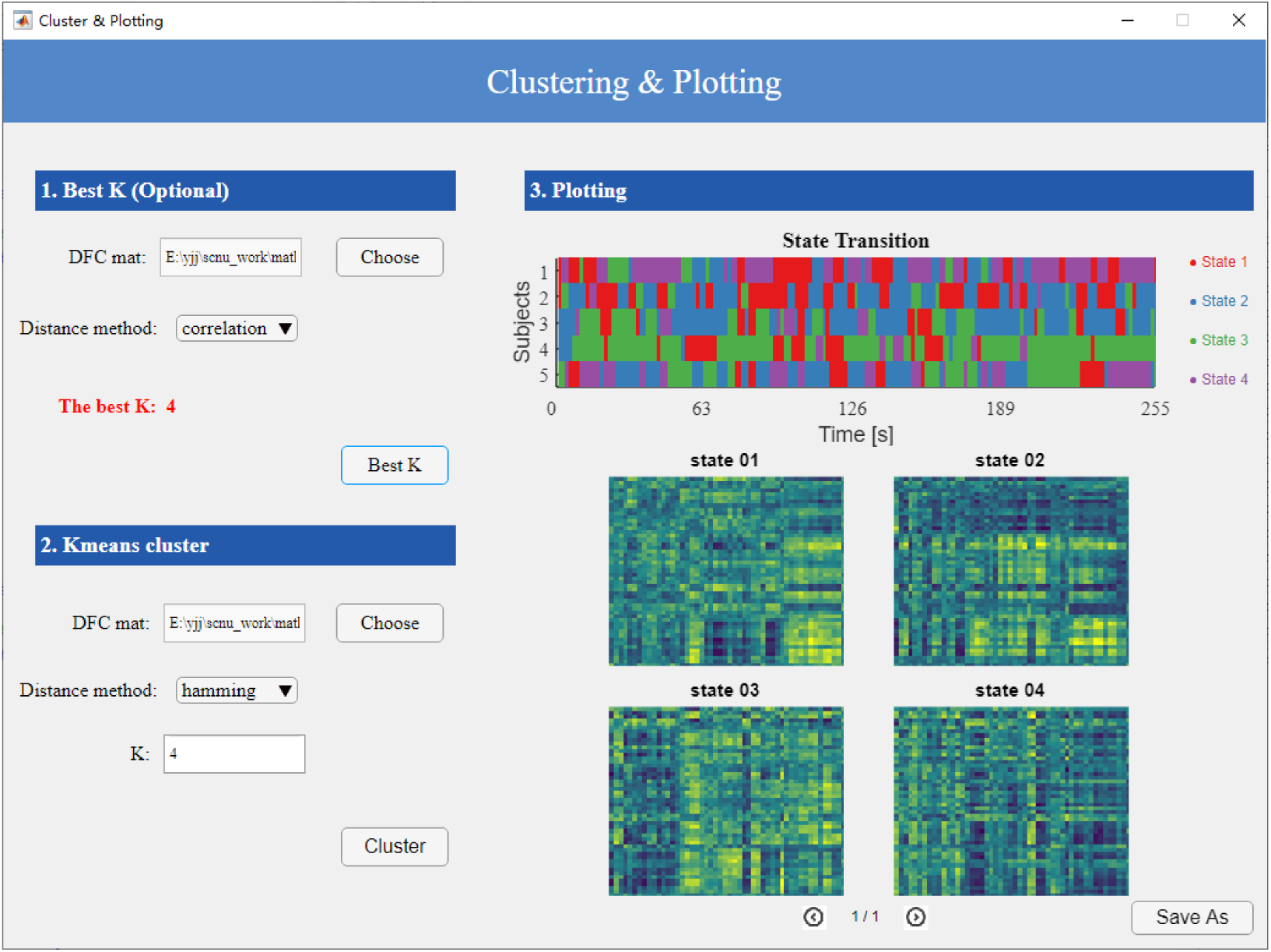
The clustering and plotting interface

In the best K-value estimation section, the user is asked to enter a file ending in "_all.mat", which serves as a summary result produced by the ROI analysis methods, and to select a clustering distance metric. The file with the extension "_all.mat" is generated as the output of the dynamic analysis methods. Clicking the "Best K" button will continue the calculation, and the interface will display the optimal K value upon completion. MATLAB’s kmeans function provides several methods for determining the distance between clusters. In this toolbox, we use "cityblock" as the default distance metric, also known as L1 distance. Users can select any distance metric based on their specific needs and the expected outcome of their analysis. Available distance measures include "sqeuclidean", "cityblock", "cosine", "correlation", "hamming".

In the K-means clustering analysis section, users can either use the Best K value obtained in the previous step or enter their desired K value. In addition, they can select a distance method for clustering and by clicking the ’Cluster’ button, the K-means clustering will be performed. This will generate K state matrices (each with dimensions [nROI, nROI]) on the right, as well as a state transition matrix (dimensions: [nSub , (nT - wsize)], where nSub is the number of subjects). However, when visualizing the state transition matrix, the x-axis is not represented by the number of time points, but by the actual time duration, calculated as (nT - wsize) multiplied by TR (time resolution). This representation allows a more intuitive understanding of the temporal significance within the state transition matrix. The results are shown in **Figure 6**.

In this module, the introduction of states and their corresponding state transition matrices enriches the depth and breadth of the analysis. The state transition matrix describes the states exhibited by all subjects at each time point, with each row representing an individual subject and each column indicating the state of that subject at a particular time point, helping researchers to understand the dynamic changes in brain networks over time. Users can also calculate relevant temporal attributes based on the state transition matrix, such as the duration of each state within specific time intervals, which helps to understand the stability of different states. Furthermore, by calculating the transition frequencies between different states, the dynamic characteristics of the brain across different states can be elucidated. Additional metrics are available in Bolton et al.(2020). This allows researchers to identify the trajectories of changes in brain activity during specific tasks or stimuli. This not only provides a more nuanced view of brain function, but also a new theoretical framework for neuroscience research.

## 3 Mathematical principles and efficacy of metrics

### 3.1 Mathematical description, formulas, free parameters, and limitation

The supplementary materials contain the mathematical descriptions and formulae for each method. Note that the mathematical principles of the extended versions are the same as those of the original versions; the only difference is that the input for the extended versions is the stimulus-evoked signals rather than the raw signals.

Table 1 summarizes the free parameters for each method. These free parameters have a significant impact on the performance and reliability of each method. Due to space limitations, we will not discuss the influence of these parameters on the results here; readers can refer to the literature cited in Table 1. In NaDyNet, we have provided the optimal free parameters for each method based on previous studies. However, it is crucial to note that these so-called best free parameters are often derived based on simulation data. Simulation data have their own limitations (Yuan et al., 2024).

In addition to the influence of free parameters, the algorithms themselves have inherent limitations. For example, the MTD algorithm requires the calculation of first-order time derivatives for two time series, which can result in a high-pass filtering effect (Shine et al., 2015). FLS takes into account two types of estimation error simultaneously (i.e. the time series values themselves and the regression coefficients), making it insensitive to data that are uncorrelated or where the covariance remains unchanged over long periods (Kalaba & Tesfatsion, 1989; Binke Yuan et al., 2024). In SWFC, the window size is often smaller than the number of ROIs, resulting in an overestimation of the estimated parameters relative to the sample size. Consequently, most current SWFC methods incorporate graphical LASSO (i.e. L1 regularization), assuming that the correlation matrix is sparse (Allen et al., 2014). In addition, dFC methods are sensitive to noise. (Binke Yuan et al., 2024), particularly framewise methods (Shine et al., 2015; Xie et al., 2019). To reduce noise sensitivity, some studies have applied sliding window averaging to the estimated dynamic network matrices, thereby increasing sensitivity to signals and improving the reliability of results (Shine et al., 2015; Xie et al., 2019). For framewise methods in NaDyNet, we do not offer a version with sliding window averaging; users are encouraged to process the data according to their specific needs.

In addition to the algorithm itself and its free parameters, a number of factors can influence algorithm efficiency, such as the preprocessing steps for fMRI data (Hutchison et al., 2013), data type (simulated vs. real fMRI-BOLD data, simulated non-fMRI-BOLD vs. simulated fMRI-BOLD data) (Shappell et al., 2019; Binke Yuan et al., 2024), SNR (Shappell et al., 2019; Binke Yuan et al., 2024), and additional steps to enhance SNR, sensitivity, and reliability (e.g., adopting inter-subject correlation and sliding window averaging). The impact of these factors on method efficacy has been thoroughly discussed in the studies listed above and will not be reiterated here.

### 3.2 Sample results to illustrate the benefits of the enhanced versions

#### 3.2.1 The logic of ISC and ISFC

The calculations for ISFC and ISC are similar to those for SFC, with the main difference being that the two time series come from different subjects (**Figure 7**). The difference between ISFC and ISC is whether the time series are extracted from the same ROI in two subjects. ISC specifically assesses the functional synchrony of two time series from the same ROI in two different subjects, whereas ISFC does not require this condition.

**Figure 7.**
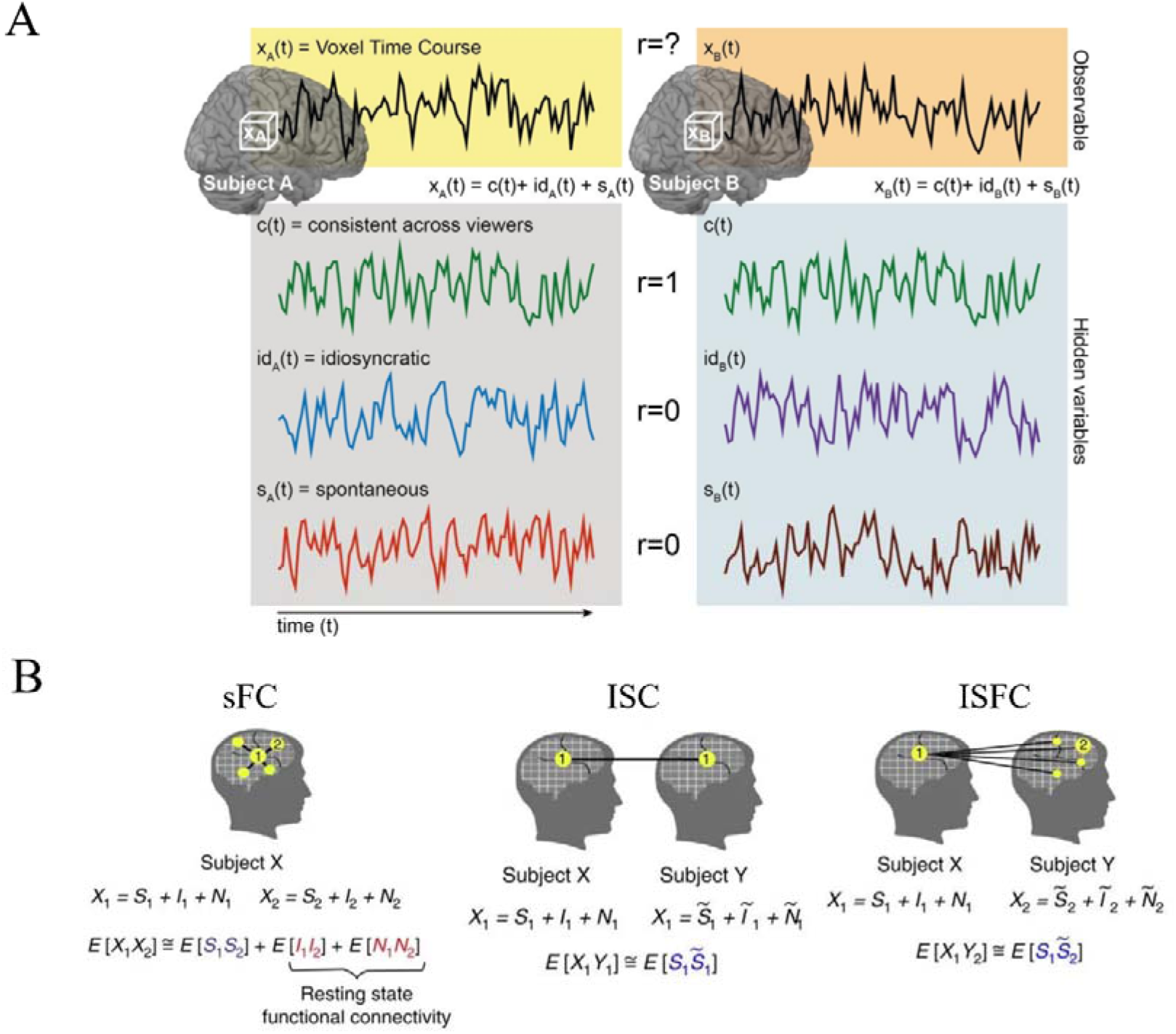
Schematic principles of SFC, ISC, and ISFC. A: Components in naturalistic fMRI signals. Adapted from Nastase et al. (2019). B: Component-dependent functional connectivity, where X1 and X2 denote two fMRI time series, and S, I, and N represent subject-shared stimulus-induced signals, subject-specific spontaneous signals, and subject-specific noise, respectively. Adapted from Simony et al. (2016).

For naturalistic fMRI data, the raw signals are regarded as a composite of three key elements: a stimulus-evoked component ( s_i_(t) ), subject-specific spontaneous fluctuations (spon_i_(t)) and non-neuronal signals (n_i_ (t)). That is: x_i_(t) = s_i_(t) + i_i_(t) + n_i_(t). The signals in s_i_(t) were shared across subjects who were presented with the same stimulus, while the latter were uncorrelated across subjects. Thus, sFC were estimates of correlations for the raw signals, whereas ISC and ISFC were estimates of correlations for the stimulus-evoked signals s_i_(t).

#### 3.2.2 ISC VS SWISC

Calculating ISC within the sliding window yields the dynamic activation results, i.e. SWISC (**Figure 8**). Dynamic activation, like dynamic functional connectivity, helps to localize cognitive and behavioral changes in real time. Song et al. (2021) focused on changes in attentional engagement during film viewing and story listening. By correlating SWISC dynamic activation results with participants’ behavioral ratings of engagement during these stimulus conditions, they found that dynamic activation of the DMN was significantly related to participants’ attentional engagement.

**Figure 8.**
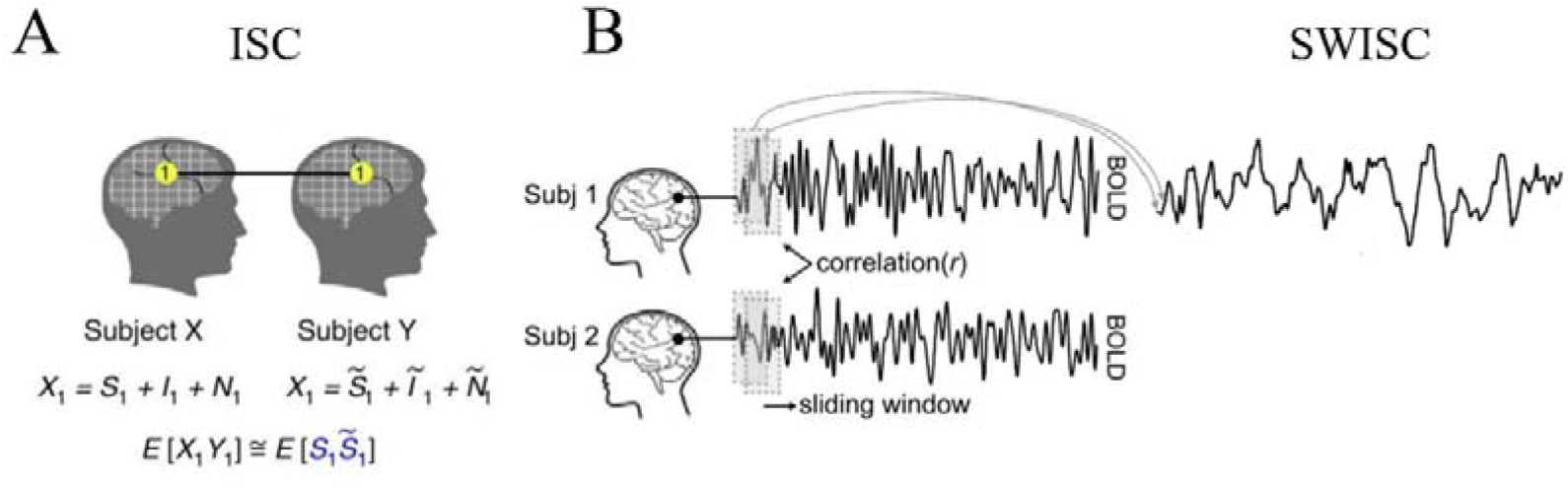
Schematic principles of SWISC. A: adapted from Simony et al. (2016). B: adapted from Song et al. (2021).

#### 3.2.2 FC vs ISFC

The default mode network is not only an intrinsic network, which is highly active in resting state (Fox & Raichle, 2007), but also a sense-making network that is critical for the comprehension of naturalistic stimuli (Yeshurun et al., 2021). Simony et al. (2016) investigated how the DMN is involved in narrative comprehension by analyzing the functional connectivity patterns in four different conditions: listening to an intact narrative, listening to a temporally scrambled versions of the narrative (word scrambled and paragraph scrambled), and resting-state with eyes open. SFC analysis found that the connectivity patterns were similar across the four conditions, whereas ISFC analysis found that the connectivity patterns within the DMN were modulated by the temporal coherence of the stimulus. Strong connectivity within the DMN was only observed in the intact and paragraph scrambled conditions (**Figure 9**). As shown in **Figure 9**, in naturalistic fMRI data, the SFC reflects the correlation between spontaneous activity signals as well as the correlation of task-evoked signals, whereas the ISFC is only sensitive to the latter.

**Figure 9.**
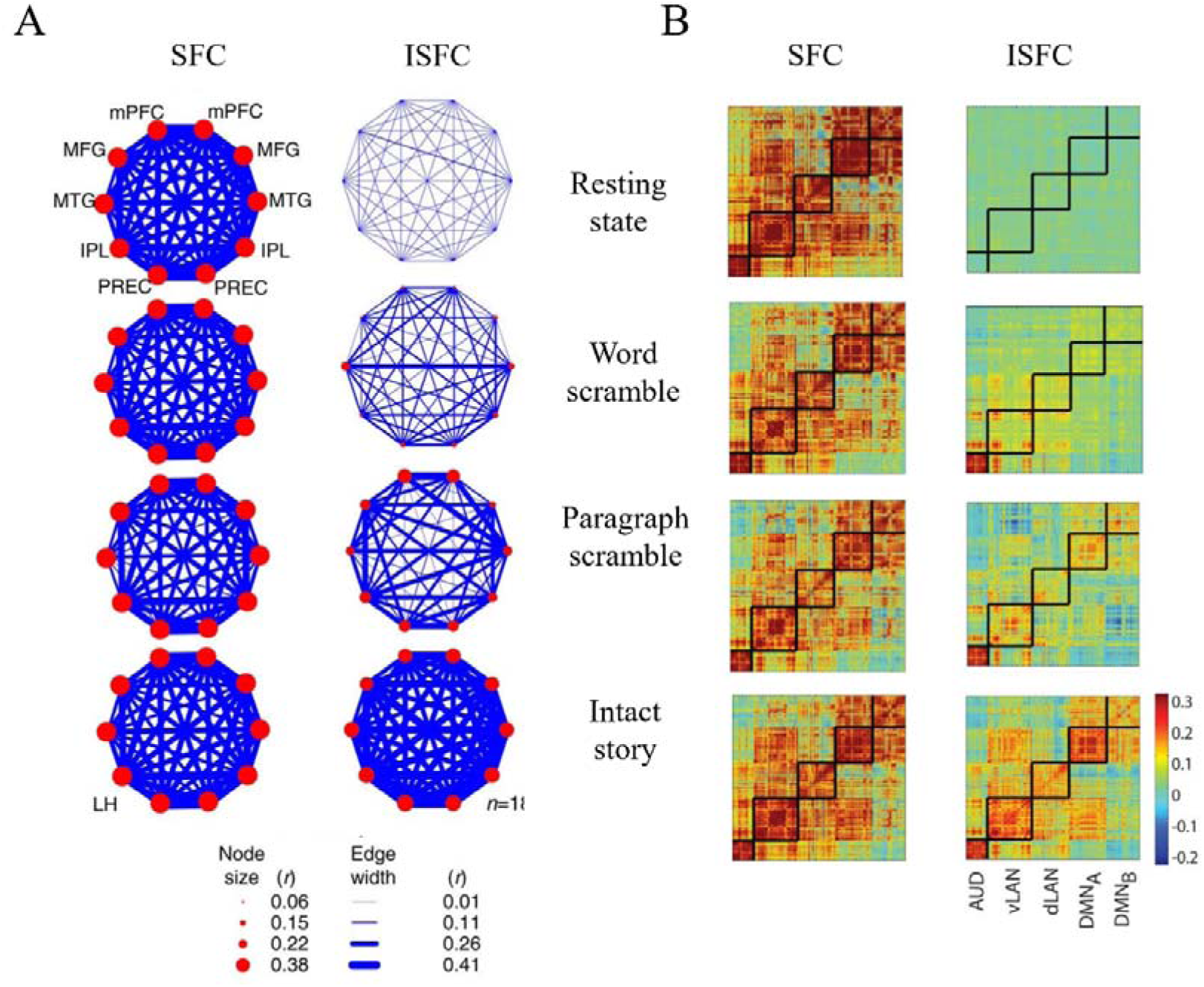
FC and ISFC results in four task conditions. A: The mean SFC and ISFC of DMN in four conditions. The ISFC was sensitive to the involvement of the DMN in narrative comprehension of the intact story. B: The mean connectivity matrices of five networks in four conditions. AUD, auditory network; vLAN, ventral language network; dLAN, dorsal language network. Adapted from Simony et al. (2016).

#### 3.2.3 SWFC vs ISSWFC

Simony et al. (2016) used SWFC and ISSWFC to compare dynamic functional connectivity patterns in intact narratives, word-scrambled narratives, and resting states. As shown in **Figure 10**, ISSWFC was more sensitive to the temporal coherence of the stimulus than SWFC. In the intact condition, the overall connectivity strength was not consistently high compared to SWFC, while the connectivity strength remained consistently low in both the word-scrambled and resting conditions.

**Figure 10.**
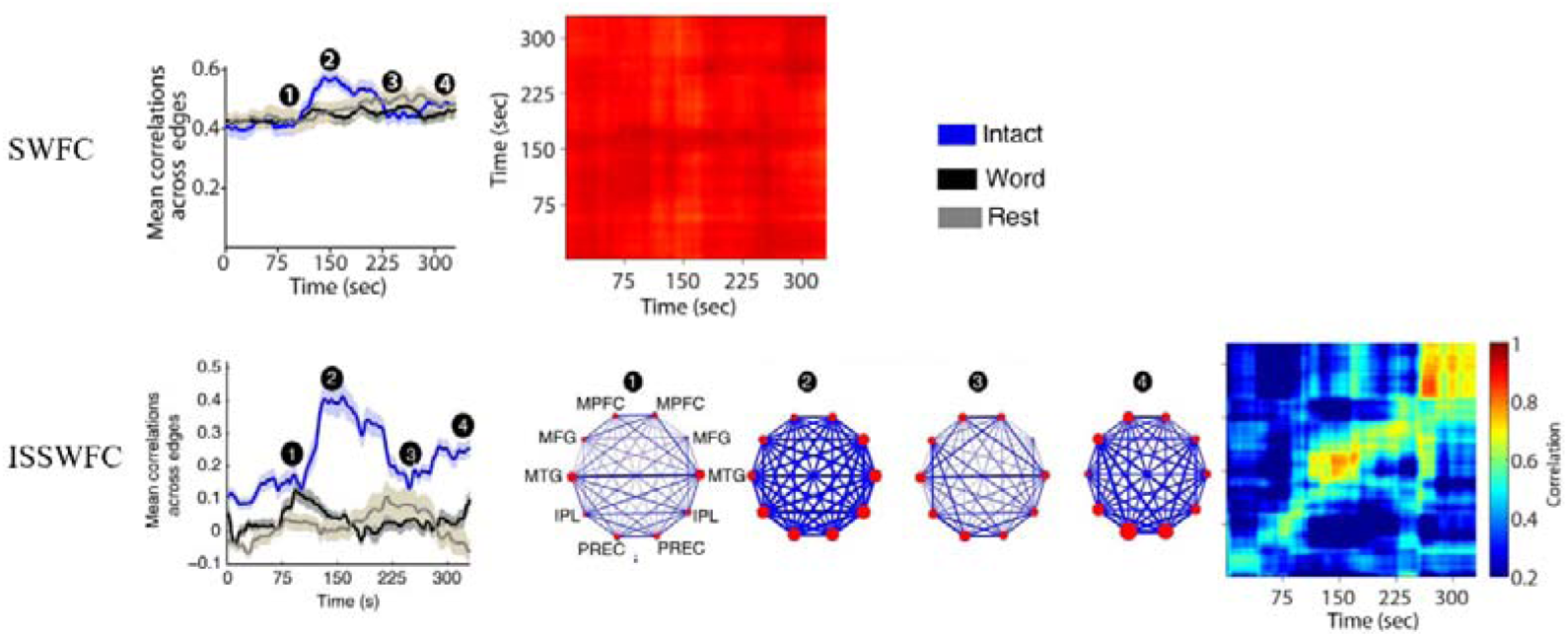
SWFC and ISSWFC results in three task conditions. top: Mean SWFC in each window and correlation matrices of all SWFC matrices over time. High mean connectivity strength was observed across the four time points for all the three task conditions. B: The mean ISSWFC in each window, the mean matrices in the four time points, and the correlation matric of all SWFC matrices over time. ISSWFC was able to track the dynamic involvement of DMN for narrative comprehension for the intact story. Adapted from Simony et al. (2016).

#### 3.2.4 DCC vs ISDCC

Chen et al. (2024) proposed ISDCC instead of DCC for naturalistic fMRI. As with ISSWFC, ISDCC used ISC to remove intrinsic and non-neuronal signals, leaving only inter-subject, stimulus-induced signals. They found that compared to DCC, ISDCC accurately revealed the underlying network reconfiguration patterns in the simulation data, identified synchronized network reconfiguration patterns across subjects, and effectively discriminated stimulus types with different temporal coherence (Figure 11). (**Figure 11**).

**Figure 11.**
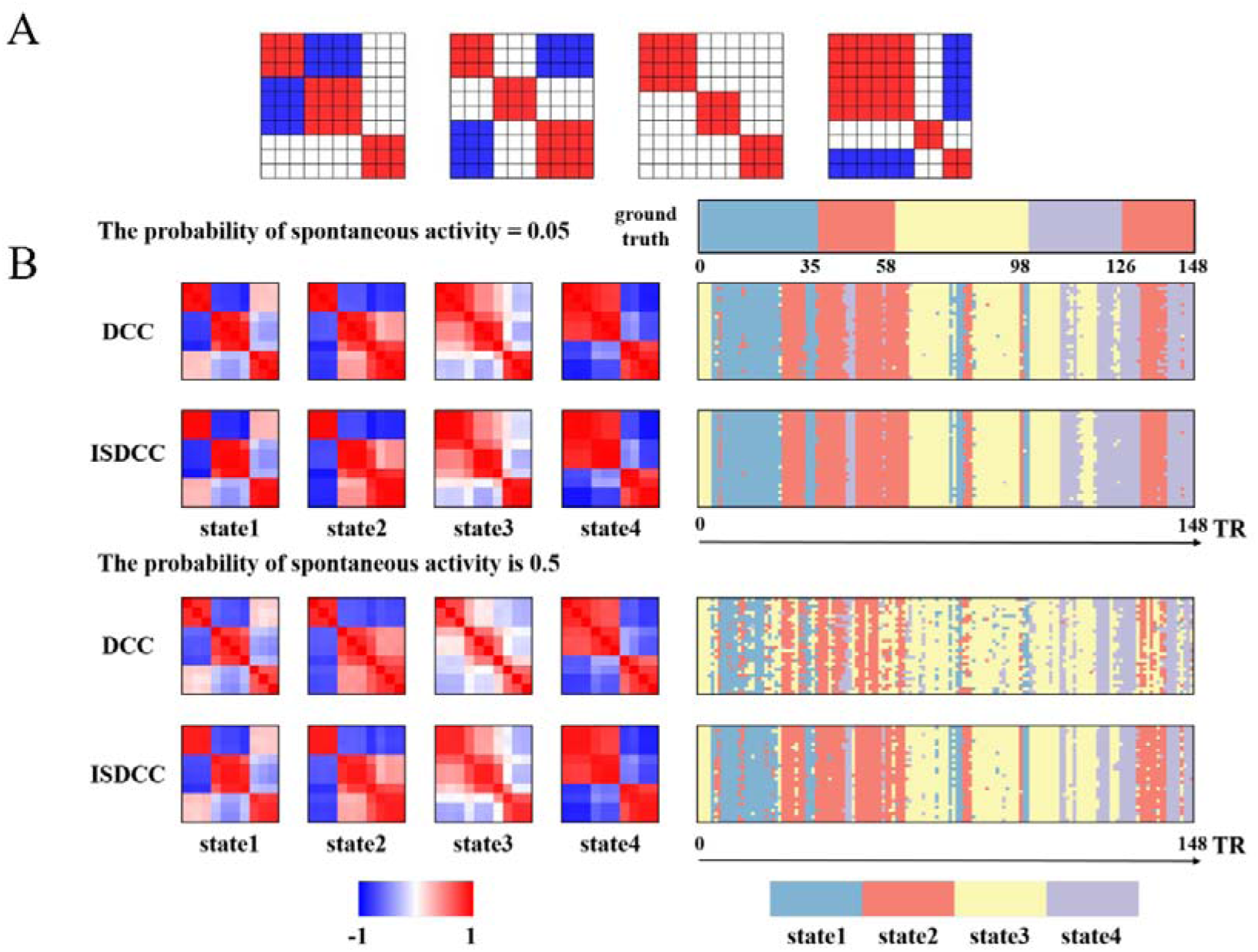
Efficacy of DCC and ISDCC assessed using simulated fMRI data. A. The predefined functional connectivity states between ten ROIs. B: The temporally reoccurring states and state transition vectors unveiled by DCC and ISDCC. The simulated fMRI signals with data-consistent signals, spontaneous signals and Gaussian noise. The simulation was repeated 30 times with a probability of spontaneous activity of 0.05 or 0.5 at each time point. Adapted from Chen et al. (2024).

These comparisons clearly show that improvements based on ISC increase the sensitivity of the methods to task-related signals. Spontaneous and task-irrelevant signals dominate natural stimulus fMRI data. Functional connectivity results based on raw signals may mislead researchers to draw incorrect conclusions.

### 3.3 Metrics efficacy

Table 2 summarizes recent advances in comparative studies of the efficacy of dFC methods. Among the six studies, SWFC and DCC are the most commonly used methods, appearing in six and four studies, respectively. In terms of data type, most studies used real fMRI data, while three used simulated data, two of which relied entirely on simulations. Efficiency metrics included test-retest reliability, differentiation of task conditions, and spatial and temporal similarity to gold standard results or to dFC metrics themselves.

**Table 2.**
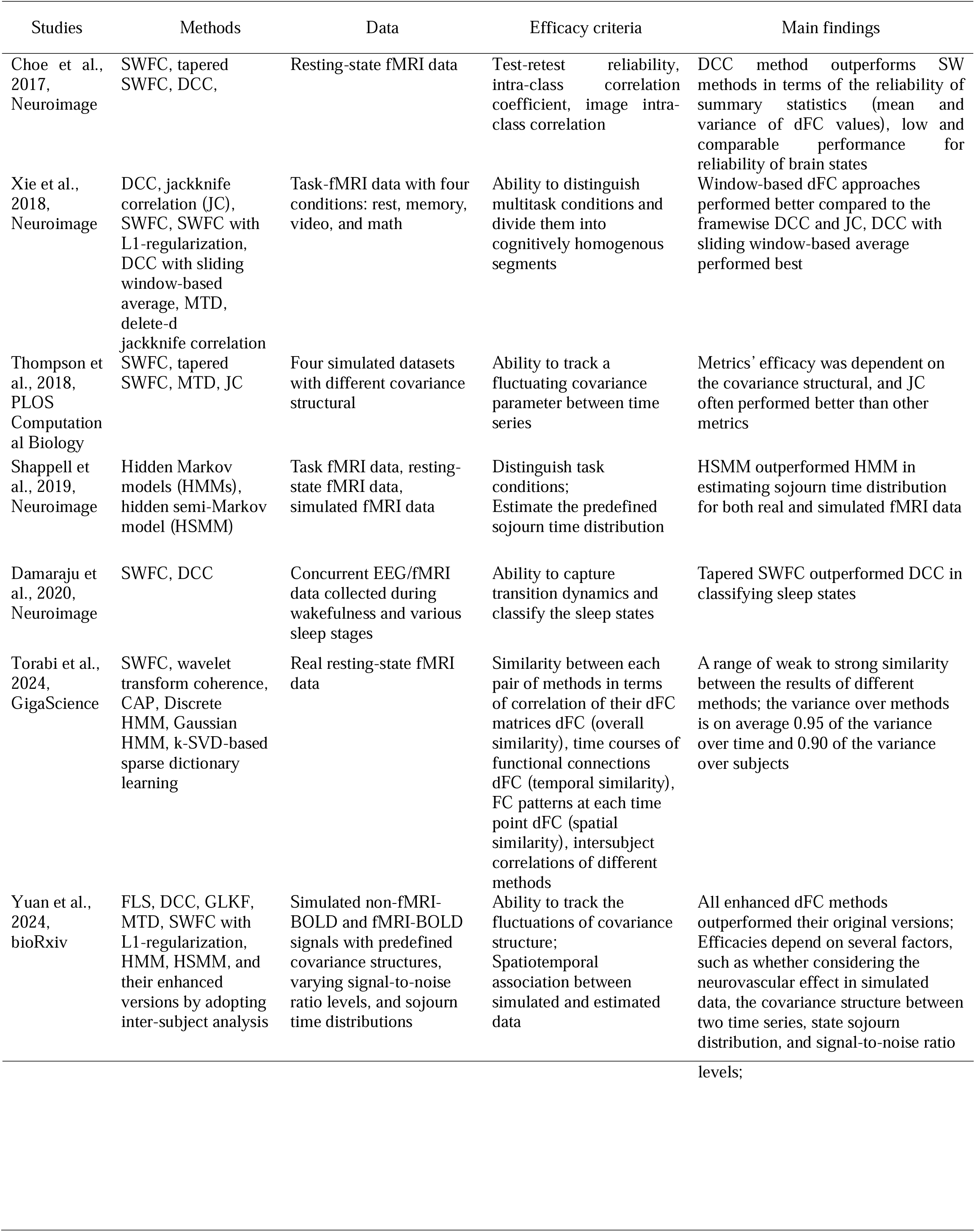
Studies comparing the efficacy of different dFC metrics

Each study focuses on different dFC methods, making it difficult to compare results across studies. For the two most commonly studied methods, SWFC and DCC, findings on their effectiveness vary between studies. For example, SWFC is better than DCC at distinguishing between different task conditions. DCC is sensitive to state transitions, often revealing multiple states within the same task or condition (Xie et al., 2018; Damaraju et al., 2020). However, in simulated data, particularly non-fMRI data (Yuan et al., 2024; Lindquist et al., 2014), DCC outperforms SWFC in tracking the rapid state transitions.

## 4 Discussion

To support research in naturalistic neuroscience, we have developed an open source Matlab-based toolbox with GUI, NaDyNet. This study provides a detailed description of the functional modules and usage of NaDyNet. NaDyNet provides six dynamic brain network analysis methods and their extended versions, as well as three static analysis methods. We presented the mathematical principles of each method, their free parameters and their limitations. We discussed the components of naturalistic fMRI data, explained the rationale for introducing ISC to improve existing dynamic network methods, and provided numerous examples to visually compare the effectiveness of the original version with the ISC-based enhanced version. In addition, we summarize the current state of research on comparing the effectiveness of dynamic network methods. However, the methods focused on in these studies vary, the selection of appropriate dynamic network methods still remains challenging, highlighting the need for more comprehensive and systematic research to compare their effectiveness.

A clear conclusion is that the efficacy of dynamic network methods is influenced by a number of factors, such as the steps in fMRI preprocessing, the free parameters of each method, and user modifications (e.g., introduction of sliding window averaging or ISC). In addition, a major problem is the lack of ground truth of connectivity states and state transition vectors in real naturalistic fMRI data. Here, we present some recommendations based on previous research. First, chronological experimental design or manipulation of experimental content helps identify timing of activation. For example, Simony et al. (2016) designed intact narrative, paragraph scrambled, and word scrambled conditions based on the time receptive window of brain language processing. Second, collecting behavioral, eye movement, and other physiological data during natural stimulus were also informative for precise mapping. For instance, Song et al. (2021) collected real-time engagement ratings from participants while listening to stories and watching films. Third, multimodal neuroimaging data, such as EEG-fMRI or separately collected SEEG and fMRI data based on the same naturalistic stimulus (Keles et al., 2024; Li et al., 2022), can be gathered for multimodal study; for example, EEG data during sleep can accurately pinpoint moments of different sleep states.

It is worth noting that most existing dynamic network methods, including those provided by NaDyNet, require clustering analysis to identify temporally recurring states. Typically, the number of states identified in dynamic brain network studies ranges from 2 to 10. Most clustering algorithms aim to maximize the distance between states, resulting in typical states with distinctive functional connectivity patterns. However, the continuous variation in the signals indicates the presence of many intermediate states. Recently, researchers have developed a TDA-based Mapper analysis method (Geniesse, Sporns, Petri, & Saggar, 2019; Hasegan et al., 2023; Saggar et al., 2022; Zhang et al., 2023), which is characterized by limited clustering of the original signal, allowing the identification of both typical and intermediate states. In addition, the mapper analysis method allows analysis at the individual level.

While the introduction of ISC improves the sensitivity of dynamic network methods to task-evoked signals, the estimation of task-evoked signals requires a group of participants undergoing the same stimulus task (Nastase et al., 2019). This group-based estimation overlooks individual differences, which hinders the application of naturalistic stimulation paradigms and dynamic brain networks in clinical settings (e.g. personalized functional localization, disease diagnosis and intervention) and education. An important question for future research will be how to individualize the decomposition of naturalistic fMRI signals. One potential solution is to simultaneously collect data from participants in both resting state and natural stimulus conditions, allowing the separation of naturalistic fMRI signals into components such as spontaneous signals and task-evoked signals based on the characteristics of the resting state signals from the previous run.

## Acknowledgments

During the development of this software, we received invaluable help and support. We sincerely express our gratitude to the following researchers: FLS: Dr. Wu, G. R. DCC: Dr. Lindquist, M. A. CAP: Dr. Bolton, T. A. W. MTD: Dr. Shine, J. M. Their contributions have been instrumental in making this toolbox a reality.

## Funding

The study is supported by the National Social Science Foundation of China (No. 20&ZD296), the Key-Area Research and Development Program of Guangdong Province (No. 2019B030335001), and the National Natural Science Foundation of China (No.32100889).

## Conflict of Interest

The authors declare that they have no conflict of interest.

## Notes

### Competing Interest Statement

The authors have declared no competing interest.

## References

1. Allen, E. A., Damaraju, E., Plis, S. M., Erhardt, E. B., Eichele, T., & Calhoun, V. D. (2014). Tracking whole-brain connectivity dynamics in the resting state. Cereb Cortex, 24(3), 663–676. doi:10.1093/cercor/bhs352

2. Binder, J. R., Conant, L. L., Humphries, C. J., Fernandino, L., Simons, S. B., Aguilar, M., & Desai, R. H. (2016). Toward a brain-based componential semantic representation. Cogn Neuropsychol, 33(3-4), 130–174. doi:10.1080/02643294.2016.1147426

3. Biswal, B., Yetkin, F. Z., Haughton, V. M., & Hyde, J. S. (1995). Functional connectivity in the motor cortex of resting human brain using echo-planar MRI. Magn Reson Med, 34(4), 537–541. doi:10.1002/mrm.1910340409

4. Bolton, T. A., Freitas, L. G., Jochaut, D., Giraud, A.-L., & Van De Ville, D. (2020). Neural responses in autism during movie watching: Inter-individual response variability co-varies with symptomatology. Neuroimage, 216, 116571.

5. Bolton, T. A. W., Tuleasca, C., Wotruba, D., Rey, G., Dhanis, H., Gauthier, B., . . . Van De Ville, D. (2020). TbCAPs: A toolbox for co-activation pattern analysis. Neuroimage, 211, 116621. doi:10.1016/j.neuroimage.2020.116621

6. Chao-Gan, Y., & Yu-Feng, Z. (2010). DPARSF: A MATLAB Toolbox for "Pipeline" Data Analysis of Resting-State fMRI. Front Syst Neurosci, 4, 13. doi:10.3389/fnsys.2010.00013

7. Chen, K., Li, C., Sun, W., Tao, Y., Wang, R., Hou, W., & Liu, D. Q. (2022). Hidden Markov Modeling Reveals Prolonged "Baseline" State and Shortened Antagonistic State across the Adult Lifespan. Cereb Cortex, 32(2), 439–453. doi:10.1093/cercor/bhab220

8. Chen, L., Tan, S., Li, C., Lin, Z., Hu, X., Gu, T., . . . Gao, X. (2024). Inter-subject dynamic conditional correlation: A novel method to track the framewise network implication during naturalistic stimuli. Brain Connect(ja).

9. Chien, H. S., & Honey, C. J. (2020). Constructing and Forgetting Temporal Context in the Human Cerebral Cortex. Neuron, 106(4), 675–686 e611. doi:10.1016/j.neuron.2020.02.013

10. Choe, A. S., Nebel, M. B., Barber, A. D., Cohen, J. R., Xu, Y., Pekar, J. J., . . . Lindquist, M. A. (2017). Comparing test-retest reliability of dynamic functional connectivity methods. Neuroimage, 158, 155–175. doi:10.1016/j.neuroimage.2017.07.005

11. Cole, D. M., Smith, S. M., & Beckmann, C. F. (2010). Advances and pitfalls in the analysis and interpretation of resting-state FMRI data. Front Syst Neurosci, 4, 1459.

12. Eickhoff, S. B., Milham, M., & Vanderwal, T. (2020). Towards clinical applications of movie fMRI. Neuroimage, 217, 116860.

13. Fan, L., Li, H., Zhuo, J., Zhang, Y., Wang, J., Chen, L., . . . Jiang, T. (2016). The Human Brainnetome Atlas: A New Brain Atlas Based on Connectional Architecture. Cereb Cortex, 26(8), 3508–3526. doi:10.1093/cercor/bhw157

14. Finn, E. S. (2021). Is it time to put rest to rest? Trends Cogn Sci, 25(12), 1021–1032.

15. Finn, E. S., Glerean, E., Hasson, U., & Vanderwal, T. (2022). Naturalistic imaging: The use of ecologically valid conditions to study brain function. Neuroimage, 247, 118776.

16. Fox, M. D., & Raichle, M. E. (2007). Spontaneous fluctuations in brain activity observed with functional magnetic resonance imaging. Nat Rev Neurosci, 8(9), 700–711. doi:10.1038/nrn2201

17. Gao, J., Chen, G., Wu, J., Wang, Y., Hu, Y., Xu, T., . . . Yang, Z. (2020). Reliability map of individual differences reflected in inter-subject correlation in naturalistic imaging. Neuroimage, 223, 117277.

18. Geniesse, C., Sporns, O., Petri, G., & Saggar, M. (2019). Generating dynamical neuroimaging spatiotemporal representations (DyNeuSR) using topological data analysis. Netw Neurosci, 3(3), 763–778. doi:10.1162/netn_a_00093

19. Glasser, M. F., Coalson, T. S., Robinson, E. C., Hacker, C. D., Harwell, J., Yacoub, E., . . . Van Essen, D. C. (2016). A multi-modal parcellation of human cerebral cortex. Nature, 536(7615), 171–178. doi:10.1038/nature18933

20. Gruskin, D. C., Rosenberg, M. D., & Holmes, A. J. (2020). Relationships between depressive symptoms and brain responses during emotional movie viewing emerge in adolescence. Neuroimage, 216, 116217.

21. Hamilton, L. S., & Huth, A. G. (2020). The revolution will not be controlled: natural stimuli in speech neuroscience. Language, cognition and neuroscience, 35(5), 573–582.

22. Hasegan, D., Geniesse, C., Chowdhury, S., & Saggar, M. (2023). Deconstructing the Mapper algorithm to extract richer topological and temporal features from functional neuroimaging data. bioRxiv. doi:10.1101/2023.10.13.562304

23. Hasson, U., Nir, Y., Levy, I., Fuhrmann, G., & Malach, R. (2004). Intersubject synchronization of cortical activity during natural vision. Science, 303(5664), 1634–1640. doi:10.1126/science.1089506

24. Hutchison, R. M., Womelsdorf, T., Allen, E. A., Bandettini, P. A., Calhoun, V. D., Corbetta, M., . . . Chang, C. (2013). Dynamic functional connectivity: promise, issues, and interpretations. Neuroimage, 80, 360–378. doi:10.1016/j.neuroimage.2013.05.079

25. Huth, A. G., de Heer, W. A., Griffiths, T. L., Theunissen, F. E., & Gallant, J. L. (2016). Natural speech reveals the semantic maps that tile human cerebral cortex. Nature, 532(7600), 453-+. doi:10.1038/nature17637

26. Jaaskelainena, I. P., Sams, M., Glerean, E., & Ahveninen, J. (2021). Movies and narratives as naturalistic stimuli in neuroimaging. Neuroimage, 224. doi:ARTN 117445 10.1016/j.neuroimage.2020.117445

27. Jia, X. Z., Wang, J., Sun, H. Y., Zhang, H., Liao, W., Wang, Z., . . . Zang, Y. F. (2019). RESTplus: an improved toolkit for resting-state functional magnetic resonance imaging data processing. Sci Bull (Beijing), 64(14), 953–954. doi:10.1016/j.scib.2019.05.008

28. Kalaba, R., & Tesfatsion, L. (1989). Time-varying linear regression via flexible least squares. Computers & Mathematics with Applications, 17(8-9), 1215–1245.

29. Keles, U., Dubois, J., Le, K. J. M., Tyszka, J. M., Kahn, D. A., Reed, C. M., . . . Rutishauser, U. (2024). Multimodal single-neuron, intracranial EEG, and fMRI brain responses during movie watching in human patients. Sci Data, 11(1), 214. doi:10.1038/s41597-024-03029-1

30. Kousta, S.-T., Vinson, D. P., & Vigliocco, G. (2009). Emotion words, regardless of polarity, have a processing advantage over neutral words. Cognition, 112(3), 473–481.

31. Li, J., Bhattasali, S., Zhang, S., Franzluebbers, B., Luh, W. M., Spreng, R. N., . . . Hale, J. (2022). Le Petit Prince multilingual naturalistic fMRI corpus. Sci Data, 9(1), 530. doi:10.1038/s41597-022-01625-7

32. Liao, W., Wu, G. R., Xu, Q., Ji, G. J., Zhang, Z., Zang, Y. F., & Lu, G. (2014). DynamicBC: a MATLAB toolbox for dynamic brain connectome analysis. Brain Connect, 4(10), 780–790. doi:10.1089/brain.2014.0253

33. Lin, N., Zhang, X., Wang, X., & Wang, S. (2024). The organization of the semantic network as reflected by the neural correlates of six semantic dimensions. Brain Lang, 250, 105388.

34. Lindquist, M. A., Xu, Y., Nebel, M. B., & Caffo, B. S. (2014). Evaluating dynamic bivariate correlations in resting-state fMRI: a comparison study and a new approach. Neuroimage, 101, 531–546. doi:10.1016/j.neuroimage.2014.06.052

35. Liu, X., & Duyn, J. H. (2013). Time-varying functional network information extracted from brief instances of spontaneous brain activity. Proc Natl Acad Sci U S A, 110(11), 4392–4397. doi:10.1073/pnas.1216856110

36. Liu, X., Zhang, N., Chang, C., & Duyn, J. H. (2018). Co-activation patterns in resting-state fMRI signals. Neuroimage, 180(Pt B), 485–494. doi:10.1016/j.neuroimage.2018.01.041

37. Lurie, D. J., Kessler, D., Bassett, D. S., Betzel, R. F., Breakspear, M., Kheilholz, S., . . . McIntosh, A. R. (2020). Questions and controversies in the study of time-varying functional connectivity in resting fMRI. Netw Neurosci, 4(1), 30–69.

38. Milde, T., Leistritz, L., Astolfi, L., Miltner, W. H., Weiss, T., Babiloni, F., & Witte, H. (2010). A new Kalman filter approach for the estimation of high-dimensional time-variant multivariate AR models and its application in analysis of laser-evoked brain potentials. Neuroimage, 50(3), 960–969. doi:10.1016/j.neuroimage.2009.12.110

39. Murphy, K., & Fox, M. D. (2017). Towards a consensus regarding global signal regression for resting state functional connectivity MRI. Neuroimage, 154, 169–173. doi:10.1016/j.neuroimage.2016.11.052

40. Nastase, S. A., Gazzola, V., Hasson, U., & Keysers, C. (2019). Measuring shared responses across subjects using intersubject correlation. Soc Cogn Affect Neurosci, 14(6), 667–685.

41. doi:10.1093/scan/nsz037

42. Pascucci, D., Rubega, M., & Plomp, G. (2020). Modeling time-varying brain networks with a self-tuning optimized Kalman filter. PLoS Comput Biol, 16(8), e1007566. doi:10.1371/journal.pcbi.1007566

43. Saarimaki, H. (2021). Naturalistic Stimuli in Affective Neuroimaging: A Review. Front Hum Neurosci, 15, 675068. doi:10.3389/fnhum.2021.675068

44. Saggar, M., Shine, J. M., Liegeois, R., Dosenbach, N. U. F., & Fair, D. (2022). Precision dynamical mapping using topological data analysis reveals a hub-like transition state at rest. Nat Commun, 13(1), 4791. doi:10.1038/s41467-022-32381-2

45. Salmi, J., Metwaly, M., Tohka, J., Alho, K., Leppämäki, S., Tani, P., Laine, M. (2020). ADHD desynchronizes brain activity during watching a distracted multi-talker conversation. Neuroimage, 216, 116352.

46. Shakil, S., Lee, C. H., & Keilholz, S. D. (2016). Evaluation of sliding window correlation performance for characterizing dynamic functional connectivity and brain states. Neuroimage, 133, 111–128. doi:10.1016/j.neuroimage.2016.02.074

47. Shappell, H., Caffo, B. S., Pekar, J. J., & Lindquist, M. A. (2019). Improved state change estimation in dynamic functional connectivity using hidden semi-Markov models. Neuroimage, 191, 243–257. doi:10.1016/j.neuroimage.2019.02.013

48. Shine, J. M., Koyejo, O., Bell, P. T., Gorgolewski, K. J., Gilat, M., & Poldrack, R. A. (2015). Estimation of dynamic functional connectivity using Multiplication of Temporal Derivatives. Neuroimage, 122, 399–407. doi:10.1016/j.neuroimage.2015.07.064

49. Simony, E., Honey, C. J., Chen, J., Lositsky, O., Yeshurun, Y., Wiesel, A., & Hasson, U. (2016). Dynamic reconfiguration of the default mode network during narrative comprehension. Nat Commun, 7, 12141. doi:10.1038/ncomms12141

50. Song, H., Finn, E. S., & Rosenberg, M. D. (2021). Neural signatures of attentional engagement during narratives and its consequences for event memory. Proc Natl Acad Sci U S A, 118(33). doi:10.1073/pnas.2021905118

51. Song, H., Park, B. Y., Park, H., & Shim, W. M. (2021). Cognitive and Neural State Dynamics of Narrative Comprehension. J Neurosci, 41(43), 8972–8990. doi:10.1523/JNEUROSCI.0037-21.2021

52. Sonkusare, S., Breakspear, M., & Guo, C. (2019). Naturalistic Stimuli in Neuroscience: Critically Acclaimed. Trends Cogn Sci, 23(8), 699–714. doi:10.1016/j.tics.2019.05.004

53. Torabi, M., Mitsis, G. D., & Poline, J.-B. (2024). On the variability of dynamic functional connectivity assessment methods. Gigascience, 13, giae009.

54. Vine, V., Boyd, R. L., & Pennebaker, J. W. (2020). Natural emotion vocabularies as windows on distress and well-being. Nat Commun, 11(1), 4525.

55. Wang, J., Ren, Y., Hu, X., Nguyen, V. T., Guo, L., Han, J., & Guo, C. C. (2017). Test-retest reliability of functional connectivity networks during naturalistic fMRI paradigms. Hum Brain Mapp, 38(4), 2226–2241. doi:10.1002/hbm.23517

56. Wang, J., Wang, X., Xia, M., Liao, X., Evans, A., & He, Y. (2015). GRETNA: a graph theoretical network analysis toolbox for imaging connectomics. Front Hum Neurosci, 9, 386. doi:10.3389/fnhum.2015.00386

57. Wang, S., Zhang, Y., Shi, W., Zhang, G., Zhang, J., Lin, N., & Zong, C. (2023). A large dataset of semantic ratings and its computational extension. Sci Data, 10(1), 106. doi:10.1038/s41597-023-01995-6

58. Willems, R. M., Nastase, S. A., & Milivojevic, B. (2020). Narratives for Neuroscience. Trends Neurosci, 43(5), 271–273. doi:10.1016/j.tins.2020.03.003

59. Xie, H., Zheng, C. Y., Handwerker, D. A., Bandettini, P. A., Calhoun, V. D., Mitra, S., & Gonzalez-Castillo, J. (2019). Efficacy of different dynamic functional connectivity methods to capture cognitively relevant information. Neuroimage, 188, 502–514. doi:10.1016/j.neuroimage.2018.12.037

60. Yan, C. G., Wang, X. D., Zuo, X. N., & Zang, Y. F. (2016). DPABI: Data Processing & Analysis for (Resting-State) Brain Imaging. Neuroinformatics, 14(3), 339–351. doi:10.1007/s12021-016-9299-4

61. Yang, Z., Wu, J., Xu, L., Deng, Z., Tang, Y., Gao, J., Li, C. (2020). Individualized psychiatric imaging based on inter-subject neural synchronization in movie watching. Neuroimage, 216, 116227.

62. Yao, S., Rigolo, L., Yang, F., Vangel, M. G., Wang, H., Golby, A. J., Tie, Y. (2022). Movie-watching fMRI for presurgical language mapping in patients with brain tumour. J Neurol Neurosurg Psychiatry, 93(2), 220–221. doi:10.1136/jnnp-2020-325738

63. Yeo, B. T., Krienen, F. M., Sepulcre, J., Sabuncu, M. R., Lashkari, D., Hollinshead, M., Buckner, R. L. (2011). The organization of the human cerebral cortex estimated by intrinsic functional connectivity. J Neurophysiol, 106(3), 1125–1165. doi:10.1152/jn.00338.2011

64. Yeshurun, Y., Nguyen, M., & Hasson, U. (2021). The default mode network: where the idiosyncratic self meets the shared social world. Nat Rev Neurosci, 22(3), 181–192. doi:10.1038/s41583-020-00420-w

65. Yuan, B., Xie, H., Gong, F., Zhang, N., Xu, Y., Zhang, H., . . . Yan, J. (2023). Dynamic network reorganization underlying neuroplasticity: the deficits-severity-related language network dynamics in patients with left hemispheric gliomas involving language network. Cereb Cortex. doi:10.1093/cercor/bhad113

66. Yuan, B., Xie, H., Wang, Z., Xu, Y., Zhang, H., Liu, J., . . . Wu, J. (2023). The domain-separation language network dynamics in resting state support its flexible functional segregation and integration during language and speech processing. Neuroimage, 274, 120132. doi:10.1016/j.neuroimage.2023.120132

67. Yuan, B., Yang, J., Guo, X., Gao, X., Hu, Z., Li, J., . . . Li, W. (2024). A systematic evaluation of dynamic functional connectivity methods using simulation data. bioRxiv, 2024.2007. 2009.600728.

68. Zada, Z., Goldstein, A., Michelmann, S., Simony, E., Price, A., Hasenfratz, L., . . . Friedman, D. (2024). A shared model-based linguistic space for transmitting our thoughts from brain to brain in natural conversations. Neuron, 112(18), 3211–3222. e3215.

69. Zhang, M., Chowdhury, S., & Saggar, M. (2023). Temporal Mapper: Transition networks in simulated and real neural dynamics. Netw Neurosci, 7(2), 431–460. doi:10.1162/netn_a_00301

70. Zuo, X., Honey, C. J., Barense, M. D., Crombie, D., Norman, K. A., Hasson, U., & Chen, J. (2020). Temporal integration of narrative information in a hippocampal amnesic patient. Neuroimage, 213, 116658.

