## Supplementary methods for "NaDyNet: A Toolbox for Dynamic Network Analysis of Naturalistic Stimuli"

**Mathematical description and formulas**

**Static functional connectivity**

Functional connectivity is often quantified using Pearson correlation coefficient,

$SFC=\frac{\sum_{1}^{T} (x_{i}(t)-\bar{x_{i}})(y_{i}(t)-\bar{y_{i}})}{\sqrt{\sum_{1}^{T} {(x_{i}(t)-\bar{x_{i}})}^{2}}\sqrt{\sum_{1}^{T} {(y_{i}(t)-\bar{y_{i}})}^{2}}}$	(1)

Where $x_{i}(t)$ and $y_{i}(t)$ are two time series from the same subjects, and *T* is the length of time series.

**Inter-subject correlation and inter-subejct functional connectivity**

ISFC and ISC also adopted Pearson correlation coefficient to estimate the Pearson correlation coefficient of two time series from two different subjects.

**Sliding-window functional connectivity with L1-regularization (SWFC)**

Given two windowed time series $x_{i}(t)$ and $y_{i}(t)$, the SWFC is:

$SWFC=\frac{\sum_{T-wl/2}^{T+wl/2} (x_{i}(t)-\bar{x_{i}})(y_{i}(t)-\bar{y_{i}})}{\sqrt{\sum_{T-wl/2}^{T+wl/2} {(x_{i}(t)-\bar{x_{i}})}^{2}}\sqrt{\sum_{T-wl/2}^{T+wl/2} {(y_{i}(t)-\bar{y_{i}})}^{2}}}$	(2)

Where$\bar{x_{i}}$ and $\bar{y_{i}}$ are the sample mean of windowed time series, and $wl$ denotes the window length.

Similar to other studies, we used a Gaussian tapered window by convolving a rectangle with a Gaussian kernel ($\sigma= 3$).

The small samples per window may lead to instable parameter estimate. L1-regularization with graphical LASSO is often adopted with assumption that the estimated functional connectivity matrix is sparse. Instead of assuming correlation

matrices being sparse, graphical LASSO assumes the inverse covariance matrix $\Phi$ to be sparse. Graphical LASSO gets the sparse solution of the inverse covariance matrix $\Phi$ by maximizing the following log-likelihood function:

$L_{1}=log det \Phi-tr(s\Phi)- \lambda\left\| \Phi\right\|_{1}$	(3)

where $\det$ denotes the determinant, $\mathrm{tr}$ denotes the trace which is the sum of all elements on the main diagonal, s is the empirical covariance matrix, $\lambda$ is the regularization parameter, and $\left\| \Phi\right\|_{1}$ denotes the L1 penalty on $\Phi$. The optimal $\lambda$ is selected through cross-validation, which estimates the optimal $\lambda$ for each subject by evaluating how well the inverse covariance matrix of a training set estimated with a given $\lambda$ describes a test set from the same subject.

**Multiplication of temporal derivatives (MTD)**

We first calculate the temporal derivative (*dt*) of each time series by performing a first-order differencing:

${dt}_{x}=x_{t}{-x}_{t-1}$	(4)

The MTD score at each time point is equal to the product of the *dt_x_* and *dt_y_*, and then normalized it by dividing each *dt* by the standard deviation (*σ*) of the *dt*:

$MTD=\frac{{dt}_{x} *{dt}_{y}}{\sigma_{x}*\sigma_{y}}$	(5)

To decrease the susceptibility to noise, a simple moving average method is adopted by averaging MTD scores surrounding a point in time within a window:

$SMAMTD=\frac{1}{2\omega+1}\sum_{t-\omega}^{t-\omega} MTD$	(6)

**Dynamic conditional correlation (DCC)**

Dynamic conditional correlation (DCC) is based on the generalized autoregressive conditional heteroscedastic model (GARCH). Bollerslev (Bollerslev, 1986) proposed the GARCH model, which was a natural generalization of ARCH (Engle, 1982). The main idea of this method is to represent the conditional variance of a single time series at some moment as a linear combination of the past conditional variance and the values of the past time series. For a GARCH (*p*, *q*) process, we assume that $y_{t}$ is a univariate time series with a conditional variance.

$y_{t}\sim N(0,\delta_{t}^{2})$	(7)

where the conditional variance $\delta_{t}^{2}$ is expressed as:

$\delta_{t}^{2}=\alpha_{0}+\sum_{i=1}^{q} \alpha_{i}y_{t-i}^{2}+\sum_{i=1}^{p} \beta_{i}\delta_{t-i}^{2} {where \alpha}_{0}>0,\alpha,\beta\geq0,\alpha+\beta<1$	(8)

The parameter $\alpha_{0}$ is a constant, $\alpha_{i}$ controls the impact of the past values of the time series, and $\beta_{i}$ controls the effects of the past conditional variance of the time series. Then GARCH(1,1) can be written as:

$\delta_{t}^{2}=\alpha_{0}+\alpha y_{t-1}^{2}+\beta\delta_{t-1}^{2}$	(9)

To illustrate the DCC model, we assume that $y_{t}$ is a bivariate time series with a covariance matrix $\Sigma_{t}$. $y_{t}$ contains two univariate time series.

$y_{t}\sim N(0,\Sigma_{t})$	(10)

The DCC method consists of two steps. First, the univariate GARCH(1, 1) model is fit to the case of two univariate time series of $y_{t}$. In the first step, Eq.(11) ~ Eq.(12) are used to estimate the standard residuals $\epsilon_{t}$.

$\delta_{i,t}^{2}=\alpha_{i,0}+\alpha y_{i,t-1}^{2}+\beta\delta_{i,t-1}^{2} for i=1, 2$	(11)

$D_{t}=diag\{\delta_{1,t},\delta_{2,t}\}$	(12)

$\epsilon_{t}={D_{t}}^{-1}y_{t}$	(13)

Second, the standardized residuals $\epsilon_{t}$(Eq.(12)) were used to estimate the covariance matrix to calculate the dynamic correlation. The $\overline{Q}$ is the non-conditional covariance matrix of $\epsilon_{t}$ (Eq.(13)). The covariance matrix is estimated after a simple scaling. The upper right corner of the covariance matrix is the correlation coefficient of the two univariate time series.

$Q_{t}=\left( 1-\theta_{1}-\theta_{2} \right)\overline{Q}+\theta_{1}\epsilon_{t-1}\epsilon_{t-1}^{T}+\theta_{2}Q_{t-1} where 0<\theta_{1}+\theta_{2}<1$	(14)

$R_{t}=diag{\{Q_{t}\}}^{-1/2}Q_{t}{\{Q_{t}\}}^{-1/2}$	(15)

$\Sigma_{t}=D_{t}R_{t}D_{t}$	(16)

DCC has been proven to outperform other dynamic methods to unveil reliable dynamic functional correlations (Choe et al., 2017; Lindquist et al., 2014), and has been used to investigate the spatiotemporal properties of language network in resting state (Yuan et al., 2023b, 2023a).

**Flexible Least Squares (FLS)**

Given two windowed time series $x_{i}(t)$ and $y_{i}(t)$ , FLS supposes that the linear regression coefficients, $\beta(t)$ changes over time,

$y_{i}(t)=x_{i}(t)\beta_{t}+\varepsilon(t)$	(17)

The idea of FLS in estimating $\beta(t)$ is minimizing two types of errors, the residual measurement error and the residual dynamic error. The residual measurement error is defined as:

$r_{m}^{2}(\beta)=\sum_{t=1}^{T} {(y_{i}(t)-x_{i}(t)\beta_{t})}^{2}$	(18)

Where *T* is the length of the time series.

The residual dynamic error is defined as:

$r_{D}^{2}(\beta)=\sum_{t=1}^{T-1} {(\beta(t+1)-\beta(t))}^{T}(\beta(t+1)-\beta(t))$	(19)

To find the residual efficiency frontier, an incompatibility cost is defined as:

$C=\mu r_{D}^{2}(\beta)$+$r_{m}^{2}(\beta)$	(20)

The incompatibility cost function $C$ generalizes the goodness-of-fit criterion function for ordinary least squares estimation by permitting the coefficient vector $\beta(t)$ to vary over time.

**General linear Kalman filter**

GLKF is an estimator of a system’s state space and covariance. It can be used to reconstruct the set of linearly independent hidden variables that regulate the evolution of the system over time:

$z(t)=(t-1)z(t-1)+\varepsilon(t)$	(21)

In Eq 20, the hidden state $z(t)$ at time t has a deterministic component given by the propagation of the previous state $z(t-1)$ through a transition matrix , and a stochastic component given by the zero-mean white noise sequence $\varepsilon(t)$ of covariance *Q(t)*.

$x_{i}(t)=H(t-1)z(t)+v(t)$	(22)

In Eq 21, the observed data $x_{i}(t)$ at time t are expressed as a linear combination of the state variable x with projection measurement matrix H, in the absence of noise. The term $v(t)$ is a random white noise perturbation (zero mean, covariance *R(t)*) corrupting the measurements.

To recursively estimate the hidden state x at each time (t = t_1_,..,t_T_), the Kalman filter alternates between two steps, the prediction and the update step. In the prediction step, the state and the error covariance are extrapolated as:

$\hat{{z(t)}^{-}}=(t-1)\hat{{z(t-1)}^{+}}$	(23)

${P(t)}^{-}=(t-1){P(t-1)}^{+}{(t)}^{T}+Q(t-1)$	(24)

where $\hat{{z(t)}^{-}}$ and ${P(t)}^{-}$ are the a priori or predicted state and the error covariance at time t, based on the propagation of the previous estimated state and covariance $\hat{{z(t-1)}^{+}}$ and ${P(t-1)}^{+}$ through the transition matrix . The superscript *T* denotes matrix transposition.

In the update step, a posteriori estimates of the state and error covariance are refined

according to:

$K(t)={P(t)}^{-}{(H(t-1){P(t)}^{-}+R(t))}^{-1}$	(25)

$\hat{{z(t)}^{+}}=\hat{{z(t)}^{-}}+K(t)(x_{i}(t)-H(t)\hat{{z(t)}^{-}})$	(26)

${P(t)}^{+}=(I-K(t)H(t)){P(t)}^{-}$	(27)

where *I* is the identity matrix, and $K(t)$ is the Kalman Gain matrix reflecting the relationship between uncertainty in the prior estimate and uncertainty in the measurements. The Kalman Gain thus quantifies the relative reliability of measurements and predictions and determines which one should be given more weight during the update step.

The optimal behavior of the Kalman filter in fMRI and physiological time series is not known. The transition matrix is usually replaced by an identity matrix I, such that the state in Eq (20) follows a first-order random walk model.

GLKF derives *R* recursively from measurement innovations and approximate *Q* as a diagonal weight matrix that determines the rate of change of ${P(t)}^{-}$. $\hat{R}$ is initialized as *I [d x d*] and adaptively updated from the measurement innovations (the pre-update residuals) as:

$\sum_{r} =\frac{{(x_{i}(t)-H(t)\hat{{z(t)}^{-}})}^{T}(x_{i}(t)-H(t)\hat{{z(t)}^{-}})}{N-1}$	(28)

$\hat{R(t)}=\hat{R(t-1)+c(\sum_{r} -\hat{R(t-1)})}$	(29)

where $\sum_{r}$ is the covariance of measurement innovations, *N* is the total number of trials and c (0 < *c* < 1) is a constant across time that regulates the adaptation speed for $\hat{R(t)}$. The constant c determines the trade-off between fast adaptation and smoothness: a small c value adds inertia to the system, reducing the ability to track and to recover from dynamic changes in the true state while a large c value increases the contribution of measurements to each update and the uncertainty associated with ${P(t)}^{-}$.

$\hat{R(t)}$ is computed before the Kalman update to replace the unknown $R(t)$ in the Kalman Gain with

$K(t)={P(t)}^{-}{H(t)}^{T}{(H(t){P(t)}^{-}{H(t)}^{T}+tr(\hat{R(t)})I_{N})}^{-1}$	(30)

where $tr$ denotes the trace of a matrix and $I_{N}$ is the identity matrix [N x N].

**Co-activation pattern analysis**

Let us consider the data matrix of estimated stimulus-evoked signals, $X_{S}\in{\mathbb{\mathbb{R}}}^{V\times T}$ where *S* is the number of subjects, *V* is the number of voxels and *T* is the number of time points. The activation or deactivation of time points was defined as ${abs(X}_{s})>Thres$, where $Thres$ is the (de) activation threshold. After having selected frames to keep for each subject, K-means clustering is used to get the population- or story-level ISCAPs. An ISCAP state or a cluster represents a matrix in which the sum of the squared distance between the data points and the cluster’s centroid is at the minimum:

$\underset{\mathcal{C}}{argmin}\sum_{i=1}^{K} \sum_{s=1}^{S} \sum_{t\in{\mathcal{\mathcal{I}}}_{s}} dist(X_{s}(\cdot,t),c_{k})$	(31)

Where *K* is the number of co-activation patterns, $\mathcal{C=}\left\{ {\mathcal{\mathcal{I}}}_{1},\cdot\cdot\cdot{\mathcal{\mathcal{I}}}_{k} \right\}$ where $\mathcal{\mathcal{I}}s$ is the set of time points that satisfies ${abs(X}_{s})>Thres$ and $c_{k}$ is the spatial map for co-activation pattern *k.*
